# A Decade of Deep Learning-based Biomedical Image Segmentation

**DOI:** 10.64898/2026.04.27.721127

**Authors:** Suhao Yu, Haojin Wang, Ningsen Wang, Sicheng Chen, Juncheng Wu, Zhenlong Yuan, Tianhao Qi, Zongwei Zhou, Fei Xia, Jun Ma, Yuyin Zhou

## Abstract

Biomedical image segmentation is a fundamental problem in computational biomedicine that aims to precisely delineate anatomical and biological structures, tissue types, or pathological regions in biomedical images. Accurate segmentation is essential for interpretation, decision-making, and quantitative analysis across a wide range of biological and medical applications. Over the past decade, the field has undergone a profound paradigm shift, evolving from task-specific specialist models to universal foundation models. This review provides an in-depth analysis of the evolution, tracing how the limitations of local discriminative learning drove the transition toward transformer-based global modeling, and large-scale generative pre-training. To help navigate the diverse landscape of interaction paradigms, we introduce the first systematic taxonomy of promptable biomedical image segmentation, categorizing existing methods into six distinct types, enabling users to intuitively select appropriate prompting strategies based on visual demonstrations and quickly pinpoint relevant literature ([Prompt Type Visualization](https://suhaoyu1020.github.io/MedicalSegmentation-PromptType-Website/)). Beyond model architectures, we discuss parallel advancements in dataset development, evaluation protocols, and application-specific adaptations across radiology, pathology, and biology. Integrating these powerful foundation models with rigorous domain-specific adaptation has great potential to improve patient outcomes and healthcare efficiency. Finally, we highlight key challenges in trustworthiness and clinical integration that must be overcome to realize the potential of the next generation of biological and medical generalists.

## 1 Introduction

Biomedical image segmentation is a fundamental task that precisely partitions medical images into biologically and clinically meaningful regions, including nuclei [1], cells [2], lesions [3], organs and tissues [4, 5]. Beyond boundary detection, it supports both biomedical research and clinical practice, spanning a wide range of applications from early-stage phenotyping and genomic screening to surgical navigation, treatment response monitoring and individualized treatment planning [6, 7].

Over the past decade, biomedical segmentation was dominated by task-specific models, typified by U-Net [8] and its variants, in which each anatomy or imaging task required a dedicated architecture, as illustrated in the top panel of Fig. 1. Although these specialist models achieved strong benchmark performance, they scaled poorly to the diversity of real-world clinical imaging, limiting their broader utility across different modalities, anatomical targets, and clinical settings [9].

**Figure 1:**
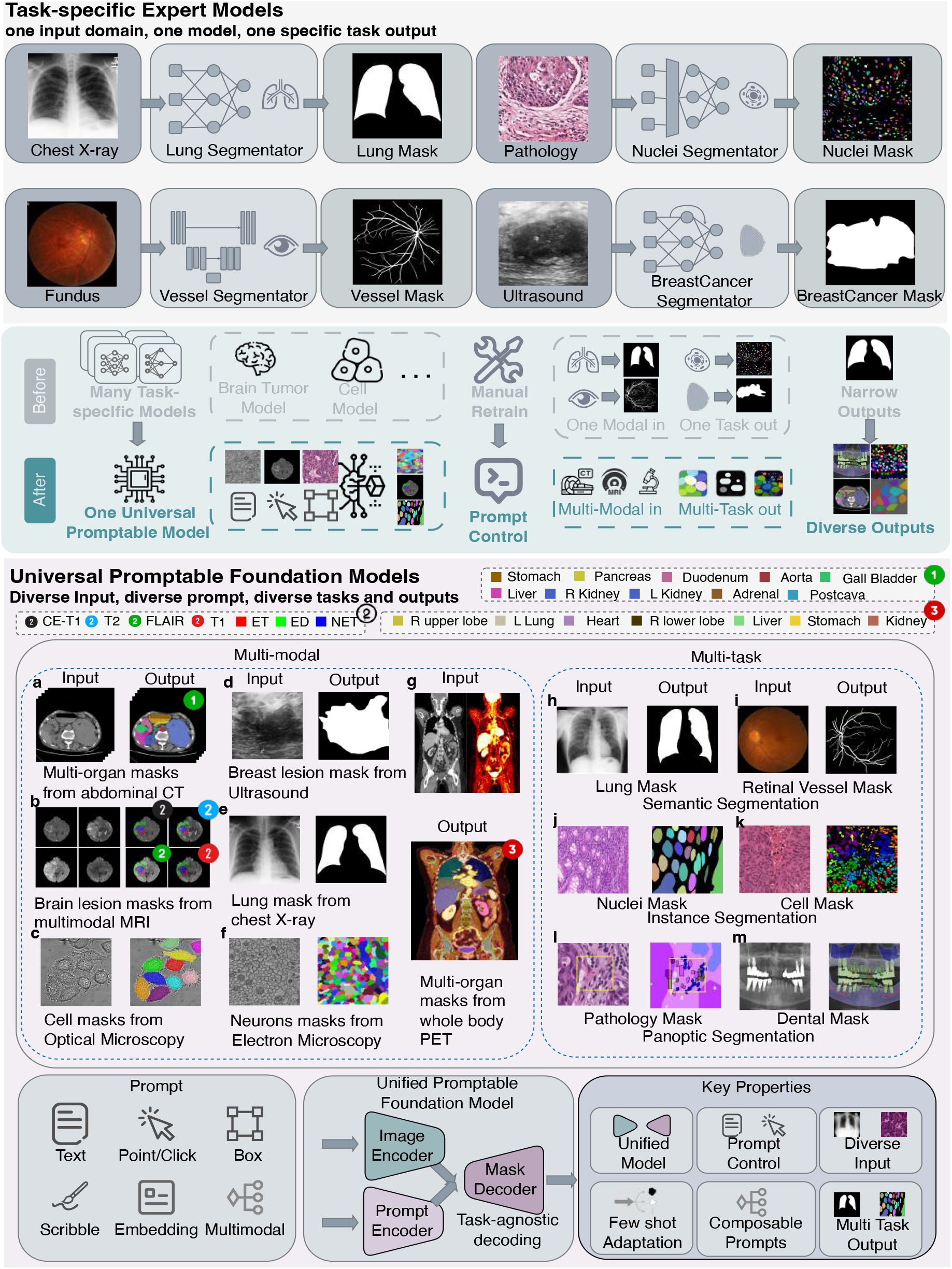
The paradigm shift from specialized experts to a universal generalist. Historically, biomedical segmentation relied on task-specific expert models (top panel), where distinct architectures were required for each anatomical target (e.g., lungs, nuclei) and imaging modality, necessitating manual retraining for every new task. In contrast, the current era is defined by universal promptable foundation models (bottom panel). These unified frameworks demonstrate broad applicability across diverse imaging modalities (e.g., CT, MRI, endoscopy) and task granularities (e.g., semantic, instance, panoptic). By using flexible user prompts such as points, boxes, or text, these models decouple task specification from model training, allowing a single frozen model to be guided without the need for retraining. Resources: **a**: AMOS [10], **b**: BRATS [3], **c**: Cell Tracking [11], **d**: BUSI [12], **e, h**: MC-CXR [13], **f**: CREMI [14], **g**: ENHANCE.PET [15], **i**: FIVES [16], **j, k**: MonuSeg-2018 [17], **l**: PanopTILs [18], **m**: Dental Panoramic Radiograph [19]

This limitation has driven a shift toward universal, promptable foundation models, exemplified by the Segment Anything Model (SAM) [20], which handle diverse anatomies, modalities, and tasks within a single model, as shown in the bottom panel of Fig. 1. This shift has been enabled by three developments: the evolution from convolutional to transformer-based architectures (see Fig. 2), large-scale pre-training on diverse biomedical datasets, and prompt-driven interaction that allows users to guide segmentation through points, boxes, or text without retraining. Models built on this paradigm, such as MedSAM [21] and CellSAM [22], now generalize across imaging applications and clinical settings, making segmentation substantially more accessible to both researchers and clinicians.

**Figure 2:**
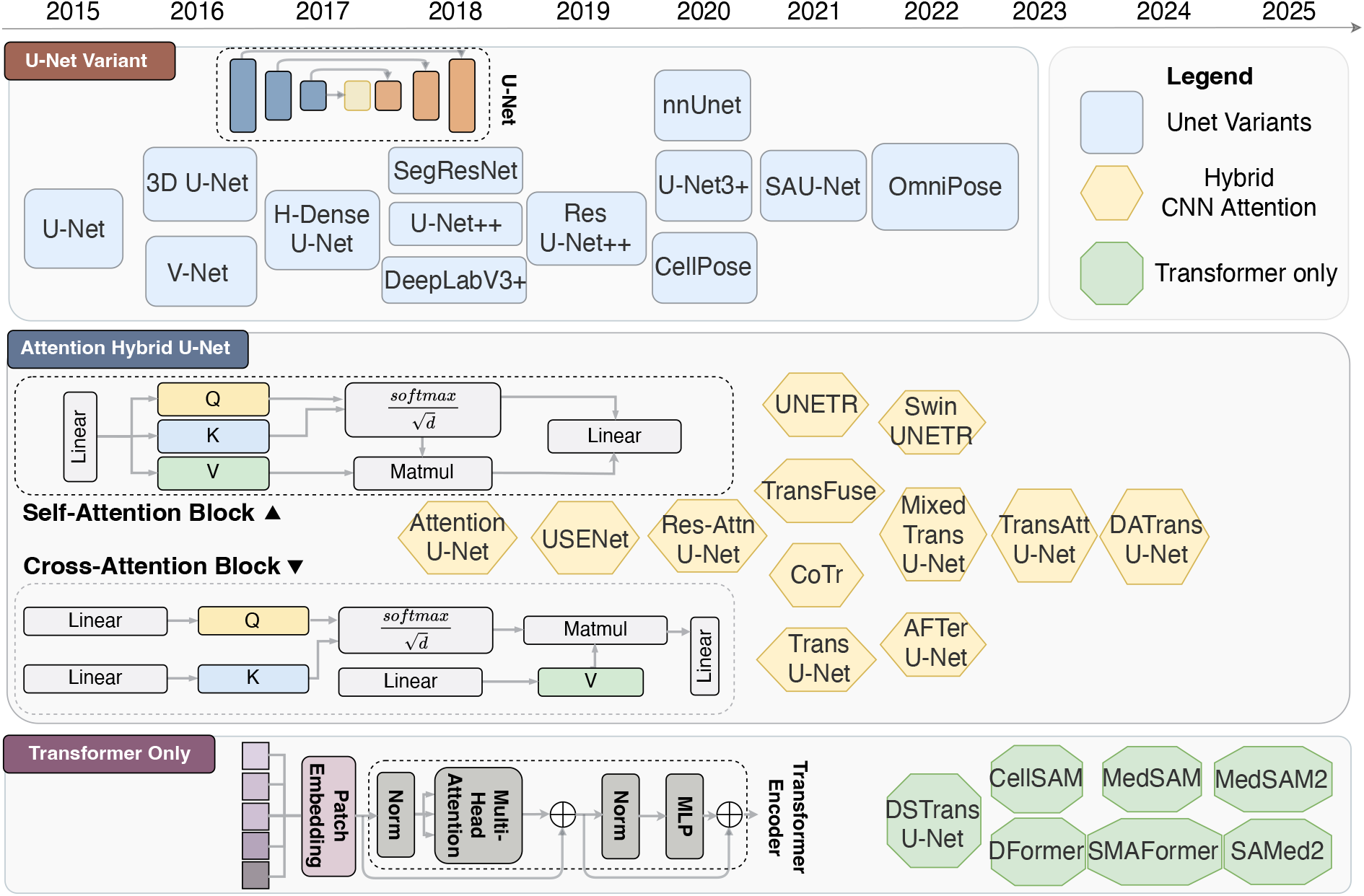
The trajectory from convolutional baselines to transformer-centric foundation models. This chronological map traces the structural evolution of segmentation networks. Early work (top row, blue) was dominated by U-Net variants, which established the encoder-decoder paradigm for modeling local image structure. As the demand for capturing broader anatomical context grew, attention-augmented and hybrid architectures (middle row, orange) emerged, combining convolutional inductive biases with attention mechanisms to integrate longer-range dependencies. The current frontier (bottom row, green) is defined by entirely Transformer-only architectures, which dispense with rigid local convolutional inductive biases and instead rely on long-range attention-based token interactions to model intricate global biological relationships, thereby providing the architectural basis for scalable promptable foundation models.

Alongside these methodological advances, segmentation has been applied across a wide range of biological and clinical settings, from organ-level analysis in radiology and neuroimaging to finer-scale tissue and cellular imaging [23]. In digital pathology, segmentation systems have reached expert-level performance in tissue analysis, enabling large-scale studies of tumor microenvironments and disease progression [24]. In biological microscopy, automated cell segmentation has become a key tool for quantitative cellular analysis, supporting high-throughput phenotypic screening across diverse experimental systems [25]. More broadly, the ability of foundation models to reduce the need for manual annotation is accelerating the adoption of segmentation tools in both research and clinical practice [26].

Despite this progress, segmentation models that perform well on benchmarks often fail to generalize across clinical settings with varying imaging protocols and patient populations [27]. Assembling the large annotated datasets needed remains a persistent challenge, as expert annotation is costly and subject to substantial variability [28]. In safety-critical settings such as surgical navigation, the inability of current models to flag uncertain predictions poses direct risks to patient safety [29]. Together, these challenges reflect a broader gap: despite successive advances from early CNNs to promptable foundation models, the field still lacks a systematic framework connecting methodological progress to reliable clinical utility [30]. Consequently, a crucial question persists: ***How can we systematically integrate these diverse algorithmic advancements to build robust, generalizable systems that meet the rigorous standards of clinical utility and scientific discovery?***

Answering this question requires a systematic re-examination of how progress in biomedical segmentation is understood and organized. Three interconnected challenges make this difficult: the rapid proliferation of promptable models has created a fragmented landscape without a clear taxonomy; adapting general vision models to biomedical data—with its volumetric structure, low contrast, and diverse modalities—remains poorly understood; and there is a persistent gap between algorithmic advances and their translation across applications, from sub-cellular microscopy to clinical radiology. Motivated by these challenges, we present a comprehensive review of biomedical image segmentation spanning a transformative decade (2015–2025), tracing the paradigm shift from task-specific specialist models to universal, promptable foundation models. We examine how architectural evolution, from convolutional networks to transformer-based models, combined with data-efficient learning strategies and prompt-driven interaction, has enabled models to generalize across the full spectrum of biomedical imaging. We further map how these systems are deployed across applications from sub-cellular microscopy to clinical radiology, and introduce a systematic taxonomy of promptable segmentation models organized by interaction modality and adaptation strategy. Together, this review provides a unified framework for understanding the rapidly evolving foundation-model landscape in computational biomedicine, and a roadmap toward more robust and clinically reliable segmentation systems.

## 2 Network Architecture Evolution: From U-Net to Transformers

Over the past decade, the architectural landscape of biomedical image segmentation has undergone a profound transformation, progressing from convolution-dominated pipelines to globally contextualized, scalable foundation models. As illustrated in the Fig. 2, a major starting point for this transition was U-Net [8], whose contracting–expanding architecture, coupled with lateral skip connections, established a canonical framework for integrating contextual semantics with precise localization. Although subsequent work refined this paradigm through improved encoder design and more effective skip connections, the inherently local nature of convolutional receptive fields increasingly constrained the modeling of complex global anatomical relationships. This limitation catalyzed the integration of attention mechanisms and, ultimately, the emergence of Transformer-only architectures. By prioritizing long-range dependency modeling over traditional inductive biases, these models paved the way for a shift from task-specific segmentation specialists to promptable foundation-model generalists capable of generalizing across biomedical domains.

### 2.1 The CNN Era

For years, U-Net [8] served as the dominant reference architecture for biomedical image segmentation, establishing a symmetric encoder-decoder framework. Building upon this foundation, several studies focused on optimizing feature aggregation and spatial recovery, which are showed in the top of Fig. 2. UNet++ [31] introduced nested, dense skip pathways to narrow the semantic gap, while ResUNet++ [32] incorporated residual connections and squeeze-and-excitation modules to improve feature representation. To further maximize full-scale information utilization, UNet 3+ [33] fused features from all network levels. Concurrently, to resolve the specific issue of blurred object boundaries, models like DeepLabv3+ [34] adapted multi-scale feature extraction to recover sharp spatial details. Further addressing this structural paradigm, to tackle the lack of robustness and interpretability in standard CNNs, SAUNet [35] introduced a parallel shape stream to capture robust geometric features alongside a dual-attention decoder for multi-level interpretability.

Parallel to these architectural refinements, the inherently three-dimensional nature of most medical scans necessitated volumetric solutions. Architectures like 3D U-Net [36] and V-Net [37] directly extended the U-Net framework into the 3D domain. However, to mitigate the prohibitive computational overhead of pure 3D architectures, hybrid models such as H-DenseUNet [38] were proposed to fuse efficient 2D intra-slice feature extraction with 3D contextual aggregation. Furthermore, addressing the scarcity of annotated 3D data, methods like SegResNet [39] coupled segmentation with image reconstruction, improving feature generalization without requiring extra labels.

Beyond structural and dimensional scaling, researchers also tackled domain-specific challenges such as cellular imaging. To overcome the poor generalization of specialized models, CellPose [25] introduced a generalist algorithm trained on highly varied datasets. Building on this, Omnipose [40] utilized distance-field information to achieve robust segmentation for cells with extreme shapes or dense arrangements.

Despite these extensive structural and domain-specific developments, the widespread adoption of nnU-Net [4] ultimately shifted the community’s attention from architectural novelty to system-level optimization. By demonstrating that carefully designed preprocessing, resampling, and training configurations could outperform many sophisticated models, nnU-Net redefined the baseline for medical image segmentation.

Even in this mature form, however, CNN-based segmentation remained fundamentally constrained by the locality of convolutional operations. Because receptive fields were built up progressively through stacked local filters, these models struggled to capture long-range anatomical dependencies and global spatial coordination, particularly in volumetric imaging and across heterogeneous organs and acquisition protocols. These limitations motivated a transition towards architectures capable of more explicit global reasoning, thereby setting the stage for the rise of Transformer-based segmentation. Detailed architectural analyses of U-Net variants are provided in Supplementary Section 4.

### 2.2 Hybrid CNN-Attention Models

A natural intermediate step towards attention-based segmentation was to augment the established U-Net framework with attention mechanisms, preserving local shape priors while capturing broader context.

As illustrated in the middle of Fig. 2, early adaptations focused on localized and channel-wise attention mechanisms to refine feature selection within convolutional architectures. For instance, Attention U-Net [41] directly embeds attention gates into skip connections to suppress irrelevant backgrounds. Similarly, USE-Net [42] employs Squeeze-and-Excitation blocks for robust feature recalibration, while Residual Attention U-Net [43] combines soft attention with residual transformations to distinguish complex disease symptoms. As the need for global context grew, hybrid Transformer-CNN architectures emerged to bridge local and long-range feature extraction. Models like TransUNet [44] (2D) and UNETR [45] (3D) replace standard convolutional encoders with Transformers to capture long-range dependencies, retaining a CNN decoder for spatial precision. Alternatively, TransFuse [46] runs Transformer and CNN branches in parallel, efficiently fusing global and local features without the redundancy of deeply stacked encoders.

Subsequent innovations tackled the computational bottlenecks and dimensional challenges of medical imaging. To mitigate the immense memory overhead of high-resolution 3D scans, CoTr [47] utilizes sparse deformable self-attention to selectively attend to informative regions, Swin UNETR [48] adopts a hierarchical approach for variable-sized tumors, and AFTer-UNet [49] balances efficiency by fusing intra and inter-slice features via axial attention. Furthermore, models like Mixed Transformer U-Net [50], TransAttUnet [51], and DA-TransUNet [52] integrate specialized modules to model cross-sample dataset correlations, self-aware non-local interactions, and dual-attention channel features, respectively.

Overall, these hybrid architectures serve as a practical compromise: they preserve the inductive biases and spatial precision of convolutions while introducing effective global context modeling. In practice, they are particularly valuable in structurally complex settings, such as multi-organ CT or brain imaging, where broad anatomical context and precise boundary delineation are equally critical.

### 2.3 From Transformer-Only Architectures to Foundation Models

Motivated by advances in large-scale representation learning, a parallel line of research progressively reduced convolutional inductive biases and explored segmentation models centered almost entirely on attention mechanisms. These Transformer-centric designs explicitly model global dependencies and long-range correlations, making them especially suitable for tasks that require extensive spatial context or multi-slice reasoning. As illustrated in the bottom of Fig. 2, representative examples include DS-TransUNet [53], which employs dual attention streams for local and global encoding. However, to mitigate the immense computational burden of standard self-attention on high-resolution medical images, subsequent architectures like D-Former [54] and SMAFormer [55] were proposed to adapt attention mechanisms on the most informative regions.

Building upon the scalability of these Transformer architectures, the field experienced a paradigm shift toward foundation-style segmentation models. Rather than optimizing a separate model for each task from scratch, these approaches seek to learn reusable segmentation capabilities through massive pre-training that can be adapted across diverse biomedical domains and imaging modalities. This shift was heavily catalyzed by the release of general-purpose vision foundation models, most notably the Segment Anything Model (SAM) [20] and its successor, SAM 2 [56]. While SAM revolutionized the field with promptable zero-shot segmentation powered by over 1 billion masks, SAM 2 extended these capabilities to continuous visual domains like video via a streaming memory architecture.

In practice, this vision has been realized most clearly through the biomedical adaptation of these models, yielding architectures like CellSAM [22] and MedSAM [21], which extend SAM to cellular and radiological imaging, respectively. Yet, because these early adaptations primarily operated in 2D, they often struggled with spatial consistency in 3D medical scans. To bridge this gap, subsequent work expanded these models to volumetric and temporal settings. For example, MedSAM2 [57] introduces memory attention to improve 3D spatiotemporal consistency, whereas SAMed-2 [58] employs a confidence-driven selective memory mechanism to mitigate noise in complex medical sequences.

## 3 Pre-Foundation Model Era: Scaling Data and Generalization

Parallel to architectural advances, biomedical image segmentation underwent a substantial transformation in learning paradigms, driven by two persistent bottlenecks: annotation scarcity and domain shift. Generating dense pixel-wise labels requires specialized expertise from radiologists and pathologists, making annotation prohibitively expensive and slow [59]. Most biomedical datasets consequently remain limited to cohorts of thousands of samples, orders of magnitude smaller than those common in general computer vision [60].

A second bottleneck lies in limited cross-site robustness. Models trained at one institution often degrade substantially when deployed at another [61]. Scanner hardware differences and imaging protocol variations are systematic sources of this degradation [62], and shifts in patient demographics compound the problem, making reliable deployment across clinical sites unlikely without explicit adaptation strategies [63].

These two challenges jointly motivated the learning paradigms illustrated in Fig. 3. To address annotation scarcity, semi-supervised, weakly supervised and self-supervised learning were developed to reduce dependence on dense expert supervision. To address domain shift, domain adaptation and federated learning were introduced to align feature representations across institutions without sharing raw patient data. The advent of Transformer-based foundation models has not replaced these paradigms but repositioned them as adaptation layers built on top of pretrained backbones: semi-supervised and self-supervised approaches for low-data fine-tuning, domain adaptation for cross-site alignment, and federated learning for privacy-preserving collaboration. We explore this transition in the following sections.

**Figure 3:**
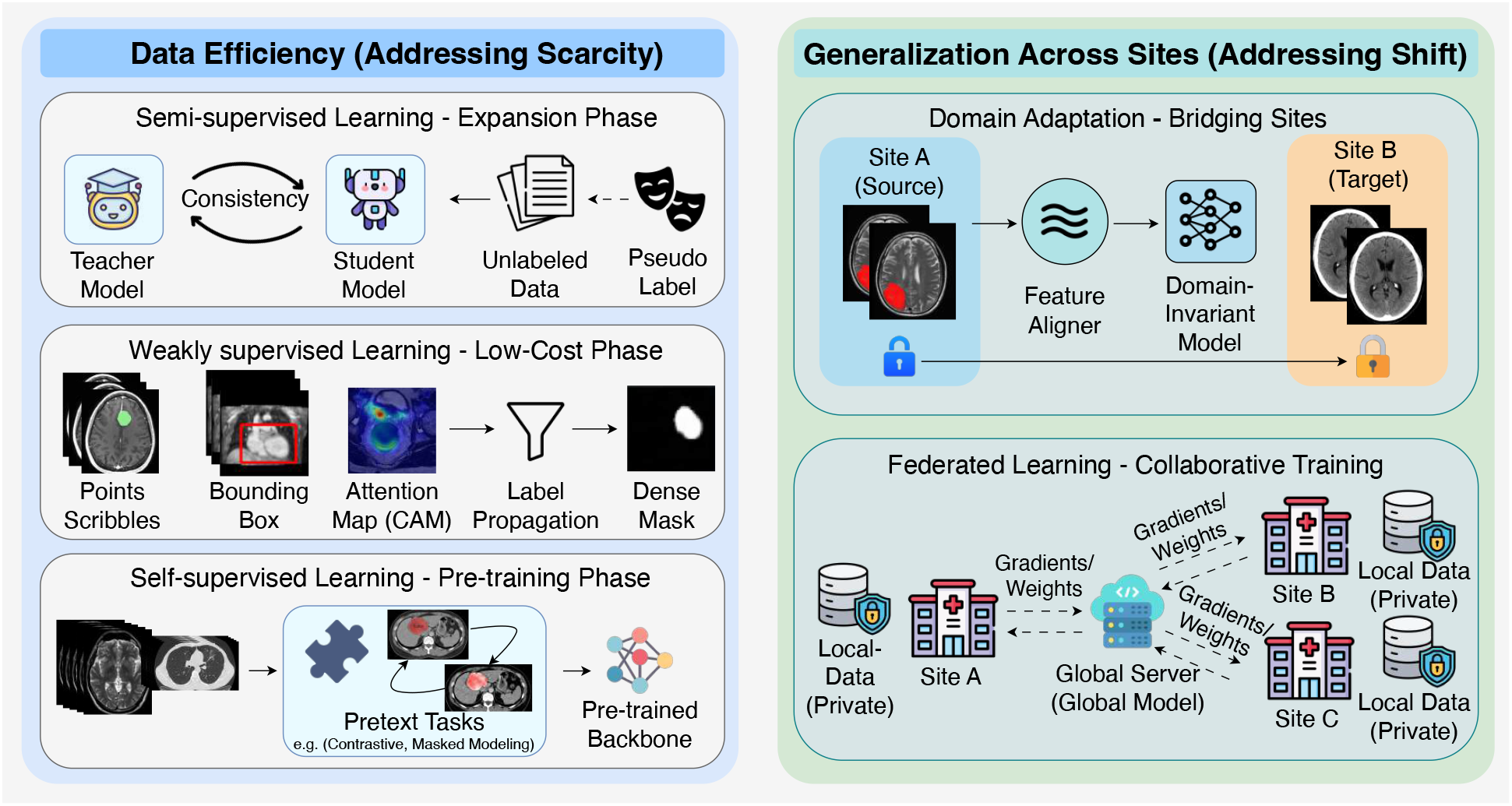
Strategic learning paradigms for addressing data scarcity and heterogeneity. Data Efficiency (Left): Semi-supervised, weakly supervised and self-supervised learning alleviate annotation scarcity through pretext tasks, the exploitation of unlabeled data, and sparse-to-dense supervision. Domain Generalization (Right): Domain adaptation and federated learning mitigate cross-site distribution shifts by aligning domain-invariant representations and enabling privacy-preserving collaborative training. Together, these paradigms laid important groundwork for the foundation model era.

### 3.1 Paradigms for Data Scarcity: Learning with Limited Labels

The high cost of expert annotation drove the development of learning paradigms that reduce dependence on dense pixel-level supervision. As left panel of Fig. 3 shows, Three strategies emerged to address this: semi-supervised learning, which pairs a small labeled set with a large pool of unlabeled images to improve performance; weakly supervised learning, which derives segmentation masks from cheaper cues such as image-level labels, bounding boxes or points; and self-supervised learning, which pretrains representations from unlabeled data using pretext tasks before transferring them to segmentation with limited labels.

Semi-supervised learning addresses the annotation bottleneck by pairing a small labeled set with a large pool of unlabeled images; most practical systems build on three complementary mechanisms: pseudo-labeling, consistency regularization and contrastive learning [64]. Cross-teaching between CNN and Transformer networks, where each model uses the other’s predictions as pseudo-labels, reduces confirmation bias by exploiting their complementary inductive biases [65]. Uncertainty-rectified pyramid consistency enforces agreement across image scales and down-weights unreliable predictions, providing more stable supervision from unlabeled data [66]. Combining global and local contrastive objectives organizes the feature space so that anatomically related regions cluster together, improving label efficiency at the pixel level [67]. Pseudo-label refinement combined with cross-modal contrastive mutual learning further reduces the performance gap when annotation quality is uneven across imaging modalities [68].

Weak supervision replaced dense pixel annotations with cheaper signals available from routine clinical or experimental practice, as illustrated by the “low-cost phase” in Fig. 3. Classifiers trained on image-level labels generated attention maps that localized regions of interest; causal refinement of these maps improved boundary accuracy by disentangling category and anatomy [69]. In histopathology, patch-level labels supported progressive multi-layer pseudo-supervision, narrowing the gap with fully annotated tissue segmentation [70]. Multiple-instance learning offered a complementary approach by treating each image as a bag of candidate regions, enabling nuclei and lesion segmentation from image-level labels alone [71]. Point annotations supported nuclei segmentation through semi-supervised detection and coarse label propagation into dense masks [72]. Scribble annotations enabled lesion segmentation via consistency-based teacher-student training, maintaining spatial structure with only sparse manual input [73]. RECIST diameter measurements, routinely recorded in clinical radiology, could be extended into full volumetric lesion masks through iterative CNN refinement and slice-wise propagation [74].

Self-supervised learning (SSL), corresponding to the “pre-training phase” in Fig. 3, sought to learn anatomy and geometry-aware representations directly from large collections of unlabeled data and then adapt them to downstream segmentation tasks using relatively few annotated examples. Unlike semi-supervised learning, which propagated scarce human labels through unlabeled images, SSL constructed its own supervision through pretext tasks and was therefore particularly useful where institutions held large but unannotated archives. Current approaches fall into three families. Contrastive methods organized the representation space by pulling together different views of the same scan while pushing apart distinct subjects; a systematic benchmark of five 3D pretext tasks [75] confirmed that such pre-training transfers reliably to downstream segmentation. Scaling this idea further, a generalizable 3D SSL framework [76] trained on 100,000 scans with multi-level image and patch contrastive objectives yielded representations that generalized across MRI and CT modalities. Reconstruction-based methods took a different route: Models Genesis [77] pre-trained 3D encoder–decoder backbones through a unified image-restoration task and transferred them across multiple organ-segmentation benchmarks. Volume-wise context restoration through 3D Rubik’s cube transformations [78] provided a simpler alternative that similarly improved generalization in pancreas and brain tissue segmentation. A third family used algorithm-generated proxy targets; supervoxel-based anomaly detection [79] enabled few-shot organ segmentation without any manual labels, and motion-based self-labeling from optical flow [80] offered a scalable variant for high-throughput cell segmentation across microscopy modalities.

The rise of foundation models has shifted the role of these paradigms. Self-supervised learning now functions primarily as a large-scale pre-training strategy that builds broadly transferable representations, rather than a standalone framework for individual datasets. Domain adaptation and federated learning have similarly moved towards lightweight fine-tuning and deployment layers that adjust shared representations to specific clinical environments without centralizing data. Progress is therefore likely to depend on integrating these roles rather than advancing each paradigm in isolation.

### 3.2 Paradigms for Generalization: Learning Across Distributed Data

Models trained on specific scanners, stains or acquisition protocols often fail when transferred to new environments, and privacy regulations frequently prevent the centralization of images needed for joint retraining. This subsection covers two paradigms that address these constraints: domain adaptation, which transfers supervision from a labeled source domain to an unlabeled target, and federated learning, which enables collaborative training across institutions without sharing patient data.

Domain adaptation (DA), shown as “bridging sites” in Fig. 3, transferred supervision from a labeled source domain to an unlabeled or sparsely labeled target domain, enabling deployment across sites without full re-annotation. Three settings emerged based on what was available at adaptation time. In *unsupervised domain adaptation* (UDA), where only unlabeled target scans were available, early methods used self-ensembling with mean-teacher networks to enforce consistent predictions across perturbed target views [81]. More recent UDA approaches combined style mixup with disentangled representation learning to separate domain-specific appearance from domain-invariant anatomy [82]. *Semi-supervised domain adaptation* (SSDA) added a small set of labeled target scans as anchors; uncertainty-aware multi-view co-training exploited these labels alongside a larger unlabeled pool to stabilize predictions under view disagreement [83]. Disentanglement with contrastive pixel-level consistency further refined SSDA by jointly separating style from content and enforcing cross-domain feature alignment [84]. *Source-free domain adaptation* (SFDA) removed access to source images entirely, relying on denoised pseudo-labels with prototype-guided uncertainty estimation to adapt pretrained models to new sites [85]. Fourier-domain style mining offered a complementary SFDA strategy by generating source-like images in the frequency domain and distilling domain-invariant features through contrastive objectives [86].

Federated learning (FL), illustrated as “collaborative training” in Fig. 3, enabled joint training across hospitals by keeping patient images on-site and sharing only model updates. Basic federated averaging, without transferring any raw scans, could match centralized performance for rare cancer boundary detection across many sites [87]. A key challenge was non-IID data, where scanner type and protocol varied across sites; personalizing both shared and site-specific network components allowed each center to correct local biases while still benefiting from pooled training [88]. Aligning local and global feature representations with contrastive objectives further improved segmentation at sites whose data differed most from the majority [89]. Many sites also had far more unlabeled scans than annotated ones; federated semi-supervised training combined pseudo-labeling and consistency regularization to exploit these unlabeled images across institutions [90]. The same strategy extended to breast MRI, where multiple perturbation strategies helped the model generalize across sites with limited annotation [91]. For large 3D architectures where communication cost was a bottleneck, split learning reduced bandwidth by transmitting intermediate features rather than full updates, with leakage-prevention mechanisms to protect patient data [92]. Multi-site feasibility studies confirmed that such schemes remained practical under realistic data imbalance and privacy constraints [93]. FL is best seen as an organizational framework for privacy-constrained collaboration, and its deployment requires governance structures alongside algorithmic design.

Despite the progress made by these paradigms, a fundamental challenge remains. Unlike natural images, biomedical structures often have ambiguous boundaries, low contrast and overlapping regions, such as adjacent soft tissues in MRI. Addressing this gap requires incorporating domain knowledge, such as anatomical constraints and imaging priors, rather than relying solely on statistical feature alignment.

## 4 Foundation Models: Towards General Biomedical Image Segmentation

Moving beyond earlier supervised and self-supervised approaches that reduced annotation demands but still required retraining per modality or target, vision foundation models (VFMs) replace task-specific architectures such as U-Net [8] with general-purpose backbones conditioned at inference time. The Segment Anything Model (SAM) [20] exemplifies this shift with a promptable interface for zero-shot segmentation. However, these models often struggle on biomedical images, where low contrast, complex shapes, and texture variation across scanners hurt pretrained 2D representations. As shown in Fig. 4 (top), generic prompts give poor results on medical data [94]. In the following sections, we review the prompt types used to guide VFMs (§4.1.1), adaptation strategies that enable reliable biomedical segmentation (§4.1.2), and the direct training of foundation models on biomedical data (§4.2).

**Figure 4:**
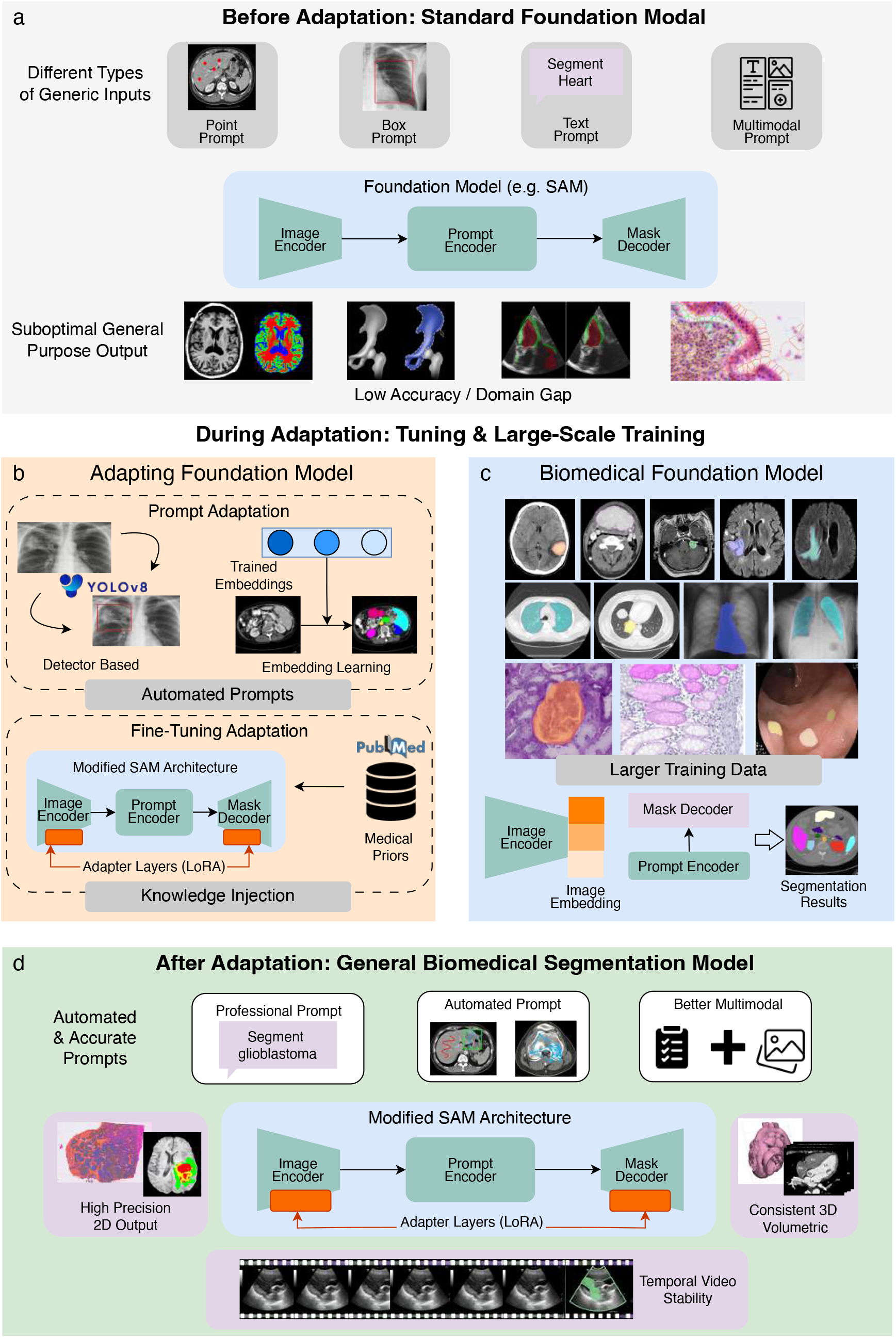
The ecosystem of prompt-guided biomedical segmentation. Before adaptation (Fig. 4a), standard foundation models perform zero-shot segmentation using generic prompts, often with suboptimal performance because of severe domain mismatch. During adaptation, two complementary pathways are used to bridge this gap. The first adapts foundation models through prompt-based methods (§4.1.1, Fig. 4b) and model-level fine-tuning (§4.1.2, Fig. 4b). The second focuses on training biomedical foundation models (§4.2, Fig. 4c) on large-scale domain-specific data to improve robustness and generalization. After adaptation (Fig. 4d), biomedical segmentation models can support specialized and automated prompts, enabling more reliable segmentation across diverse modalities and tasks.

### 4.1 Adaptation of Foundation Models

#### 4.1.1 Adaptation via Prompting

Promptable segmentation, popularized by the Segment Anything Model (SAM) [20], recasts segmentation as a prompt-conditioned inference problem, letting users steer a general-purpose model toward specific targets without retraining. Because biomedical tasks span different scales and modalities, no single prompt suffices, we therefore organize this section around six prompt types (Fig. 4): point, box, and mask prompts for spatial guidance; text and embedding prompts for semantic guidance; and multimodal prompts that combine the two. In the following, we review these categories in turn and discuss how they help bridge the gap between VFMs and domain-specific biomedical tasks.

##### Explicit Spatial Prompts

Explicit spatial prompts localize regions of interest via geometric constraints in pixel space, typically through lightweight point or box interfaces. Point prompts use sparse clicks for coarse localization [21], while box prompts specify a rectangular extent for more stable guidance at slightly higher annotation cost [94]. Unlike the coarse geometric constraints provided by points or boxes, masks encode detailed spatial priors and are therefore well suited to irregular anatomical structures with ambiguous boundaries [95], yet such density can propagate biologically implausible structure into the final prediction [96]. Recent work has explored ways to improve mask-prompt robustness [97], but high-stakes clinical use still calls for boundary-level quality control and explicit fallback criteria.

All three spatial prompt types share a common limitation that they rely on explicit manual geometric specification which leads to inconsistent performance across different settings [21]. Interactive refinement extends spatial prompting from one-shot instruction to an iterative process by updating prompts in response to intermediate predictions [98, 99]. Automated initialization reduces reliance on manual placement by using upstream detectors such as YOLOv8 [100] to generate coarse boxes that initialize SAM [101] (Fig. 4, middle-left). At the frontier of interactive segmentation, recent work has begun to explore agentic reinforcement learning (RL), in which interaction is formulated as a sequential decision-making problem [102]. By training across diverse clinical scenarios, agentic RL may navigate ambiguous regions more robustly than fixed refinement policies to narrow the gap between human intent and model execution.

##### Semantic Prompts

Semantic prompts address the limitations of purely spatial cues by shifting the emphasis from geometric localization to conceptual specification [103]. Text prompts provide a natural-language interface through which users can specify biomedical targets using free-form descriptions to enable zero-shot segmentation across datasets [104]. However, the inconsistency of biomedical terminology makes segmentation performance strongly dependent on language–vision alignment [105]. Embedding prompts replace explicit language with learned representations reusable across experimental settings [106], yet this loss of human readability obscures why a particular segmentation was produced. Overall, text and embedding prompts trade off interpretability against reusability, making semantic prompts high-level query interfaces that rely on careful design and spatial-prompt fallbacks for difficult or safety-critical targets [104].

##### Multimodality Integrated Prompts

Multimodal prompt fusion combines language with spatial indicators, using language to clarify intent and geometry to constrain localization—especially valuable in crowded or low-contrast scenes [107]. Such complementarity is especially attractive for large-scale biomedical studies, although it also amplifies the need for consistency checks. By integrating these complementary signals, multimodal prompting can reduce repeated manual correction and improve consistency across annotators and sites [108]. At the same time, this increased adaptivity raises system complexity to design choices, especially in automated or high-throughput biomedical pipelines [109]. Effective deployment therefore benefits from standardized prompting protocols and attention to computational efficiency, as increased interaction and inference cost may become practical bottlenecks at scale.

#### 4.1.2 Adaptation via Model Fine-tuning

As discussed in §4.1.1 and illustrated in Fig. 4, prompting provides a non-invasive means of steering general-purpose models, but its reliance on external cues often falls short of capturing the full complexity of biomedical data. In biomedical segmentation, small localized errors induced by imperfect prompting can propagate through inference and introduce substantial quantitative bias [21]. These limitations motivate a shift from purely external control to intrinsic model adaptation through fine-tuning.

##### Task-Specific Fine-tuning of Foundation Models

Extensive benchmarks indicate that SAM’s zero-shot capability is often unreliable in biomedical imaging—stronger for large, visually separable targets under careful prompting [110], but degrading on low-contrast structures and volumetric data with subtle boundaries [111]. Therefore, fine-tuning is motivated less by architectural novelty than by distributional mismatch in imaging or annotation protocols. For example, *µ*SAM [112] shows that fine-tuning foundation models on microscopy datasets improves segmentation quality across light and electron microscopy modalities, and Poly-SAM [113] fine-tunes SAM for colon polyp segmentation in colonoscopy images and demonstrates that task-specific adaptation can outperform both vanilla SAM and specialized segmentation models. More recently, inspired by SAM3 [114], MedSAM-3 [103] explores fine-tuning SAM3 on semantically annotated medical data to support text-driven segmentation and further investigates agent-in-the-loop refinement with multimodal large language models, reporting strong performance across multiple imaging modalities.

##### Parameter-Efficient Adaptation

Parameter-efficient adaptation addresses the computational cost and catastrophic forgetting of full fine-tuning [115]. In high-throughput settings, adapters can standardize prompt generation and reduce operator dependence by shifting part of the interaction burden into learned, reusable modules [116, 101]. Related work explores lightweight prompt learning under weak supervision [117] and architectural modifications [118] as complementary strategies for rapid, low-overhead specialization. Taken together, these studies suggest that efficient adaptation of SAM requires preserving its promptable interface while augmenting the model with lightweight structures.

##### Extending SAM to 3D and Volumetric Segmentation

Since biomedical modalities like CT and MRI are volumetric while the original SAM is 2D, recent work adapts SAM to capture cross-slice dependencies [119]. A prominent strategy involves incorporating parameter-efficient adaptation modules [115], while other approaches [120, 121] extend promptable segmentation directly to volumetric inputs. Motivated by similar goals, MedicalSAM-2 [122] reframes both 2D and 3D medical segmentation as a video object tracking problem. Collectively, these studies show that effective 3D adaptation of SAM depends on preserving the promptable interface while augmenting the model with mechanisms that account for volumetric context. A comprehensive survey of fine-tuning strategies and architectural modifications for adapting foundation models to biomedical imaging is provided in Supplementary Section 3.

### 4.2 Training Biomedical Foundation Models from Scratch

Beyond adapting existing foundation models, a more fundamental strategy is to train specialized foundation architectures directly on large-scale biomedical datasets (Fig. 4, middle-right), replacing generic visual pre-training with biomedical-centric objectives to align representations with clinical image characteristics [21, 123]. An overview of the large-scale 2D, 3D, and video datasets that underpin foundation model pre-training and evaluation is provided in Supplementary Section 2. Parameter-efficient adaptation of natural-image VFMs such as SAM [124] is efficient but limited by the domain gap, so biomedical-native models still lead on tasks such as lesion or cell differentiation despite higher training cost [123]. MedSAM2 [57] extends foundation-model training to 3D images and medical video by incorporating domain-aligned training and large-scale interactive annotation. SAM-Med3D [120] explores training 3D foundation models directly on volumetric medical data. Because domain-specific priors are embedded directly into these models, lighter forms of downstream specialization often become more effective and stable than when applied to generic vision backbones [125, 126]. Training biomedical foundation models therefore not only improves zero-shot generalization, but also lowers the barrier to efficient downstream adaptation, offering a complementary route between full retraining and purely prompt-based control.

Collectively, these adaptation strategies and specialized training regimes mark a substantial shift towards more reliable and prompt-driven biomedical analysis. As illustrated in Fig. 4 (bottom), they allow adapted biomedical models to move beyond generic behavior and provide biologically and clinically meaningful segmentation across diverse modalities and spatial dimensions. Yet, this apparent maturity invites closer scrutiny. Despite the remarkable zero-shot capabilities of vision–language foundation models, our synthesis of the current landscape reveals an important caveat: the illusion of understanding. Many promptable models successfully segment salient regions on the basis of low-level visual cues rather than genuine anatomical comprehension. We therefore argue that trustworthy medical segmentation requires evaluation criteria that extend beyond conventional overlap-based metrics to include uncertainty-aware reasoning and anatomical coherence, thereby ensuring that models can recognize the limits of their own competence.

## 5 Applications, Challenges and Future Directions

The transition from task-specific architectures to versatile foundation models has substantially expanded the operational scope of biomedical image segmentation. Beyond improvements in benchmark performance, the true significance of this shift lies in its downstream impact. Broadly, this impact is expressed in two complementary domains: **biological research**, where segmentation accelerates discovery by quantifying cellular phenotypes and spatial interactions at scale; and **clinical medicine**, where robust delineation of organs and lesions improves diagnostic precision, treatment planning, and procedural safety. We summarize this ecosystem in Fig. 5, mapping current applications together with persistent bottlenecks and plausible future directions. This section examines how modern segmentation paradigms are reshaping these domains, while highlighting the challenges that must be addressed for segmentation to deliver biomedical insight.

**Figure 5:**
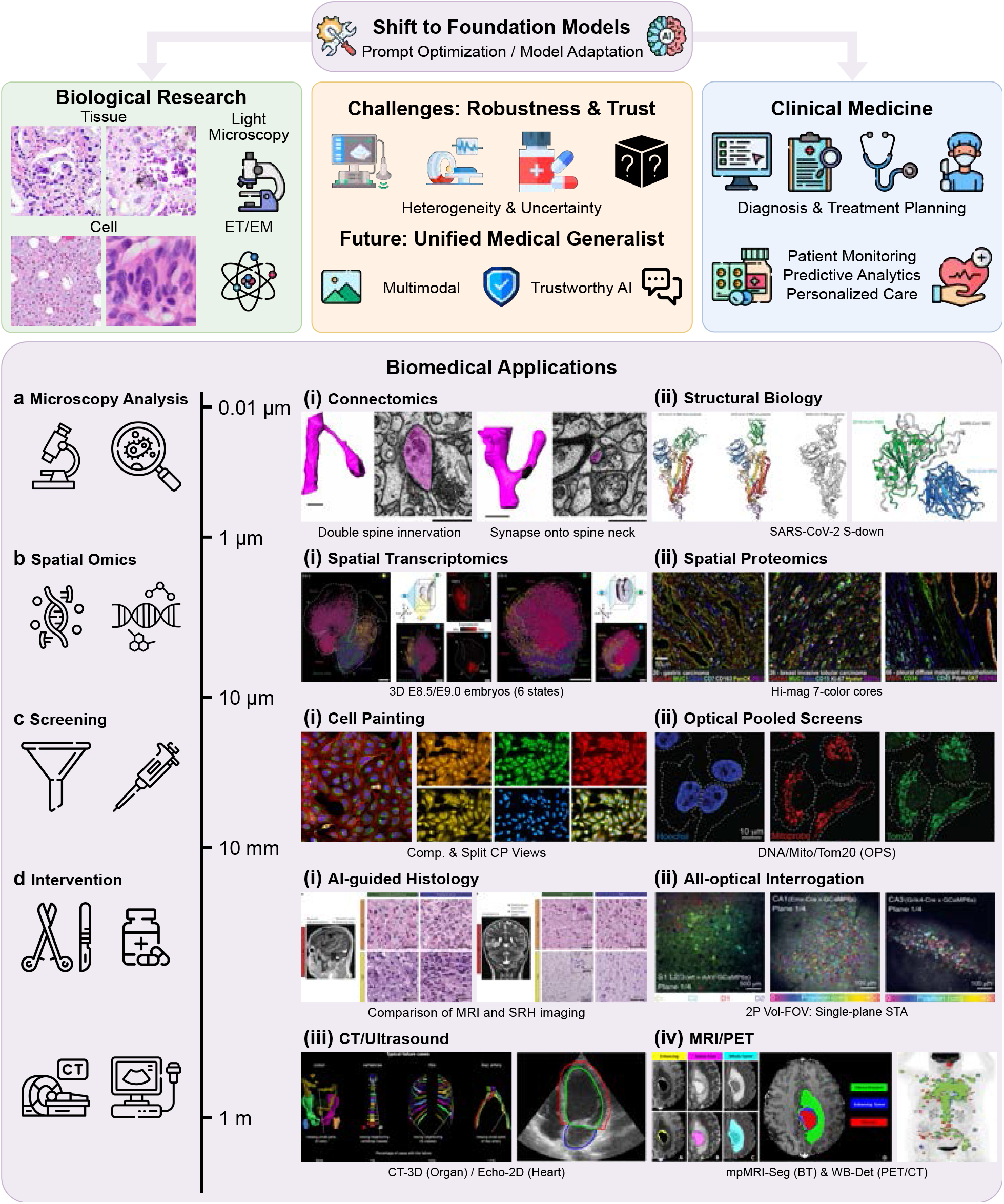
The landscape of biomedical foundation models: applications, bottlenecks, and future horizons. The diagram summarizes the operational scope and likely evolutionary trajectory of biomedical AI. **Shift to foundation models:** the upper panel illustrates the transition toward prompt-optimized models, supporting two broad domains: Biological Research, spanning tissue to near-atomic scales through microscopy and ET/EM; and Clinical Medicine, enhancing diagnosis, treatment planning, and personalized care. **Challenges and future directions:** widespread deployment remains constrained by barriers in robustness and trust, arising from heterogeneity, uncertainty, and domain mismatch, and motivating the development of a unified medical generalist capable of multimodal and trustworthy reasoning. **Biomedical applications:** the lower panel maps representative downstream tasks across spatial scales (0.01*µ*m to 1m): (a) microscopy analysis, including connectomics and structural biology; (b) spatial omics, including spatial transcriptomics and proteomics; (c) screening, including high-throughput paradigms such as Cell Painting and optical pooled screens; and (d) intervention, including clinical applications ranging from AI-guided histology and all-optical interrogation to volumetric radiology modalities such as CT, MRI, and ultrasound.

### 5.1 Accelerating Biological Discovery: From Microscopy to Spatial Omics

In the realm of biological discovery, image segmentation often forms the critical bottleneck between raw microscopy data and quantitative insight. This bottleneck has been substantially alleviated by generalist algorithms such as CellPose [25] and StarDist [127], which demonstrated that models trained on diverse biological images could generalize across microscopy modalities and approach human-level performance without task-specific tuning.

Vision foundation models have further lowered the barriers to large-scale image analysis. Microscopy-oriented adaptations of SAM, including MicroSAM [112] and CellSAM [22], have introduced stronger zero-shot generalization in complex tissue environments. By integrating SAM’s promptable interface with domain-aware components, these frameworks help bridge interactive prompting and automated screening, particularly for multiclass cell detection. In parallel, fluorescence-oriented foundation models, such as UniFMIR [128] and CytoImageNet-based approaches [129], exploit self-supervised learning on millions of unlabeled images to capture subtle subcellular dynamics.

Beyond morphology alone, segmentation has become a prerequisite for spatial omics, where accurate delineation of cellular boundaries is essential for assigning transcriptomic or proteomic measurements to individual cells [130]. Transformer-based segmentation frameworks have shown increasing ability to parse dense tissue architectures at subcellular resolution, enabling more precise mapping between gene expression and spatial location. For example, methods such as SCS [131] integrate histological imagery with sequencing spots to refine cellular boundaries in low-contrast regions. More recently, generative AI has begun to address one of the most persistent bottlenecks in basic research—severe data scarcity—by enabling robust segmentation in ultra-low-data regimes [60]. Diffusion-based augmentation methods [132], for instance, can synthesize realistic training variation from a small number of examples, allowing laboratories to deploy high-performance segmentation models even for rare phenotypes or newly developed assays.

### 5.2 Broadening the Clinical Horizon: From Screening to Intervention

In clinical medicine, segmentation has evolved far beyond simple anatomical delineation and has become a core component of quantitative precision medicine and preventive care. In radiology, fully automated whole-body segmentation systems have enabled so-called opportunistic screening, transforming routine diagnostic CT into a source of broader health assessment [23]. By automatically quantifying muscle mass, visceral fat, and bone mineral density without additional radiation exposure or acquisition cost, these models allow clinicians to identify otherwise under-reported risks such as sarcopenia and osteoporosis. In digital pathology, segmentation increasingly acts as a gateway to computational prognostics. By isolating tumour stroma, infiltrating lymphocytes, and related microenvironmental structures in gigapixel slides, AI systems can identify sub-visual morphological biomarkers that predict treatment response and patient outcomes more accurately than standard grading alone [24].

The role of segmentation becomes even more consequential when moving from diagnosis to treatment. In radiation oncology, deep learning has substantially reduced the bottleneck of contouring organs at risk, compressing treatment planning from hours to minutes while maintaining expert-level consistency [133]. In the operating room, the integration of real-time segmentation into augmented-reality guidance systems is beginning to reshape intraoperative navigation [134]. By overlaying segmented critical structures—such as vessels or nerves—directly onto endoscopic or surgical video, these systems provide a dynamic safety map that can adapt to tissue deformation and thereby help reduce perioperative complications.

### 5.3 Challenges and Future Directions

Despite the remarkable architectural evolution from U-Net to promptable foundation models, a persistent gap remains between benchmark performance and reliable utility in biomedical practice. Much of this disconnect originates not only from model design, but also from how data are curated and how evaluation is performed. Over the past decade, benchmark resources have grown considerably in both scale and diversity. Early challenges such as BraTS [3] focused on a single organ and cohorts of only hundreds of cases, whereas more recent datasets such as TotalSegmentator [23], AbdomenAtlas [135], and multimodal resources such as BiomedParse [123] contain thousands of annotated volumes spanning dozens of organ classes and modalities, from radiology to microscopy. A comprehensive taxonomy of over 60 such benchmarks is provided in Supplementary Section 1 and 2. Yet most existing benchmarks remain rooted in fixed-label, closed-set evaluation. Only a small fraction natively supports the prompt-driven interaction modes that foundation models are designed to enable. Spatially prompted resources are currently the most mature: for example, MedSAM [21] was trained on 1.57 million image–mask pairs spanning 10 imaging modalities, but such resources remain predominantly 2D, leaving volumetric prompt-based evaluation underdeveloped. Text-grounded segmentation benchmarks are even rarer: BiomedParse [123] pairs images with text descriptions and masks across nine modalities, but its scale remains limited relative to the combinatorial diversity of biomedical terminology. Benchmarks for multimodal compositional prompting, such as those combining text with boxes, are largely absent. This mismatch between the promptable paradigm and the prevailing benchmark ecosystem represents a major bottleneck. Without prompt-diverse and interaction-aware evaluation resources, the field lacks reliable signals for comparing and improving foundation models under realistic use conditions. Formal definitions of segmentation evaluation metrics across semantic, instance, and panoptic settings are provided in Supplementary Section 5. Beyond dataset design, current foundation models still largely operate as black boxes and typically lack intrinsic mechanisms for uncertainty quantification [136]. In high-stakes settings such as radiation oncology or surgery, a segmentation system must do more than produce accurate contours: it must also communicate confidence, identify ambiguous boundaries, and flag out-of-distribution inputs for expert review. Progress over the next decade will therefore depend on addressing dataset design, interaction-aware evaluation, uncertainty estimation, and clinical integration as tightly coupled problems rather than independent axes of development.

#### Architecture

U-Net-style models are likely to remain important, but increasingly as efficient spatial backbones embedded within flexible systems rather than as fixed end-to-end networks trained for closed sets of labels [8, 4]. Promptable control will probably become the dominant interaction interface, allowing users to specify intent, location, and granularity at inference time rather than hard-coding tasks and classes into the model [20, 57, 137]. This trajectory reflects clinical reality, in which questions vary across patients and a single deployed model must adapt without retraining. Interactive refinement is also likely to become an intrinsic component of segmentation systems, because many clinically important boundaries are heterogeneous or ambiguous and require iterative correction under stable model behaviour [95, 138, 135, 139]. More broadly, models must operate reliably on full 3D volumes and across time, with performance judged by volumetric and longitudinal consistency rather than slice-wise accuracy alone [140, 141].

#### Dataset

The most valuable datasets will likely evolve beyond the conventional “images with masks” format toward multimodal, longitudinal evidence packages, while retaining voxel-wise annotations where they remain reliable and clinically meaningful [142, 3]. Imaging data will increasingly be paired with radiology reports, patient context, and follow-up outcomes, because many important findings are long-tailed, difficult to enumerate in advance, and sometimes not directly annotatable in early disease stages [143, 144, 145]. Longitudinal cohorts are likely to become a major axis of future growth, as subtle disease signals often emerge as change over time and models must learn what early progression looks like relative to baseline [146]. Dense masks will remain indispensable for anatomy, therapy planning, and quantitative measurement, but will increasingly be deployed strategically rather than universally, complemented by weaker or indirect supervision when boundaries are uncertain. Dataset quality will therefore shift from local curation alone to broader governance, emphasizing coverage across institutions, scanners, and populations, together with explicit characterization of label uncertainty and missingness.

#### Task

A longer-term unifying formulation may be vision-language question answering, in which both the query and the response are multimodal and segmentation becomes one tool among several rather than the default output [147]. In this framing, classical U-Net-style segmentation becomes a conditional capability: masks are produced on demand, guided by prompts that specify anatomical intent, temporal context, or diagnostic purpose, rather than predefined label spaces. Clinical questions are inherently intent-driven and context-dependent, such as assessing suspicion, progression, longitudinal change, or supporting evidence—often conditioned on patient history [148]. Correspondingly, the outputs of these systems must be compositional and grounded, combining textual interpretation, structured measurements, and spatial evidence such as segmentations or retrieved image regions when appropriate. Reasoning in such systems will need to remain auditable [149, 150], with explicit links between claims and the image regions, measurements, or longitudinal comparisons that support them, rather than opaque end-to-end predictions.

Looking further ahead, segmentation is likely to evolve from a standalone task into a core capability within tool-augmented medical agents. Rather than relying exclusively on internal weights, foundation models may increasingly act as orchestrators that dynamically invoke external tools to verify and ground their spatial predictions. However, the deployment of agentic systems in safety-critical biomedical settings introduces serious challenges. One central concern is agency drift, in which reinforcement-learning-based agents optimize imperfect reward signals at the expense of clinical common sense, potentially producing plausible but incorrect segmentations in rare or atypical cases. Furthermore, tool augmentation also introduces the possibility of compounding error, whereby a hallucinated downstream output or a misinterpreted external knowledge source propagates through the reasoning chain. The development of robust and verifiable environments for training medical agents, together with strict guardrails for tool invocation, therefore, remains a prerequisite for meaningful clinical translation.

## 6 Conclusion

In this review, we chronologically trace the development of biomedical image segmentation from specialized task-specific models to promptable foundation models in the transformative past decade (2015-2025). We outline the fundamental paradigm shift from task-specific, convolutional expert models, exemplified by U-Net, to versatile foundation models that can be guided through prompts. This transition is driven by a broader move from local convolutional representations to globally contextualized Transformer-based architectures and large-scale pre-training. In parallel, learning paradigms such as self-supervised learning and federated learning addressed two of the field’s most persistent challenges: annotation scarcity and cross-site heterogeneity. Through prompt-based adaptation and biomedical-specific pre-training, foundation models have evolved from generic zero-shot tools to more specialized, interactive systems. Their capabilities have also expanded beyond 2D image segmentation to encompass volumetric and temporal settings, opening the way to open-vocabulary, text-guided segmentation and agent-in-the-loop refinement. Realizing the full potential of this paradigm will require models that support 3D and temporal reasoning, quantify uncertainty, and integrate transparently into real-world scientific and clinical workflows. Ultimately, the next stage of progress will depend not only on improved model capability, but also on building segmentation systems that are reliable, adaptable, and trustworthy enough to translate effectively from data to decision.

