## Supplementary Material for "A Decade of Deep Learning-based Biomedical Image Segmentation"

### A Decade of Deep Learning-based Biomedical Image Segmentation – Supplementary Information

#### 1 Biomedical Segmentation Datasets and Benchmarks

Table 1: Summary of biomedical segmentation benchmarks.

| Benchmark | Year | Modality | Segmentation Target | Segmentation Type | Scale | Link |
|---|---|---|---|---|---|---|
| BraTS | 2012<br>2013<br>2014<br>2015<br>2016<br>2017<br>2018<br>2019<br>2020<br>2021<br>2022<br>2023<br>2024<br>2025 | MR | Brain Tumor | Semantic | 35<br>35<br>200<br>200<br>200<br>331<br>351<br>460<br>494<br>1470<br>1470<br>1470<br>1538<br>N/A | https://www.synapse.org/Synapse:syn53708126/wiki/626320 |
| PROMISE | 2012 | MR | Prostate | Semantic | 50 | https://promise12.grand-challenge.org/ |
| ISLES | 2015<br>2016<br>2017<br>2018<br>2022 | MR<br>MR<br>MR<br>MR/CT<br>MR | Stroke Lesion | Semantic | 28<br>35<br>43<br>63<br>250 | https://www.isles-challenge.org/ |
| FeTS | 2021<br>2022<br>2024 | MR | Brain Tumor | Semantic | 341<br>1470<br>1470 | https://fets-ai.github.io/Challenge/ |
| FeTA | 2021<br>2022<br>2024 | MR | Fetal Brain Tissue | Semantic | 80<br>120<br>120 | https://fetachallenge.github.io/pages/About |
| AIMS-Tbi | 2024<br>2025 | MR | Traumatic Brain Injury Lesion | Semantic | 489<br>652 | https://aims-tbi.grand-challenge.org/<br>https://aims-tbi25.grand-challenge.org/ |
| ATLAS R1.1 | 2017 | MR | Stroke Lesion | Semantic | 220 | https://fcon_1000.projects.nitrc.org/indi/retro/atlas_R1.html |
| ATLAS R2.0 | 2022 | MR | Stroke Lesion | Semantic | 955 | https://atlas.grand-challenge.org/ |

*Continued on next page*

| Benchmark | Year | Modality | Segmentation Target | Segmentation Type | Scale | Link |
|---|---|---|---|---|---|---|
| MAMA-MIA | 2025 | MR | Primary Breast Tumor | Semantic | 1564 | https://www.ub.edu/mama-mia/ |
| LiTS | 2017 | CT | Liver Tumor | Semantic | 131 | https://competitions.codalab.org/competitions/17094 |
| KiTS | 2019<br>2021<br>2023 | CT | Kidney Tumor | Semantic | 210<br>300<br>489 | https://kits19.grand-challenge.org/<br>https://kits-challenge.org/kits21/<br>https://kits-challenge.org/kits23/ |
| COVID-19 Lung CT Lesion Segmentation Challenge 2020 | 2020 | CT | COVID-19 lung lesion | Semantic | 199 | https://covid-segmentation.grand-challenge.org/COVID-19-20/ |
| AutoPET | 2022<br>2023<br>2024<br>2025 | FDG-PET/CT | Whole Body Tumor | Semantic | 1014<br>1014<br>1611<br>1611 | https://autopet.grand-challenge.org/<br>https://autopet-ii.grand-challenge.org/<br>https://autopet-iii.grand-challenge.org/<br>https://autopet-iv.grand-challenge.org/ |
| HECKTOR | 2020<br>2021<br>2022<br>2025 | FDG-PET/CT | Head & Neck Tumor | Semantic | 201<br>224<br>524<br>≥1200 | https://www.aicrowd.com/challenges/miccai-2020-hecktor<br>https://www.aicrowd.com/challenges/miccai-2021-hecktor<br>https://hecktor.grand-challenge.org/<br>https://hecktor25.grand-challenge.org/ |
| ToothFairy1 | 2023 | CBCT | Inferior Alveolar Canal | Semantic | 480 | https://toothfairy.grand-challenge.org/ |
| ToothFairy2 | 2024 | CBCT | Multi-structure dental segmentation(42 classes) | Semantic | 480 | https://toothfairy2.grand-challenge.org/ |
| ToothFairy3 | 2025 | CBCT | Multi-structure dental segmentation(77 classes) | Semantic | 532 | https://toothfairy3.grand-challenge.org/ |
| VerSe | 2019<br>2020 | CT | Spine | Semantic | 120<br>300 | https://github.com/anjany/verse |
| JSRT-Seg1<br>JSRT-Seg2 | 2000 | X-ray | Lung Field<br>Lung Field & Heart | Semantic | 60<br>247 | http://db.jsrt.or.jp/eng.php |
| SIIM-ACR | 2019 | X-ray | Pneumothorax | Semantic | 12047 | https://www.kaggle.com/c/siim-acr-pneumothorax-segmentation |
| MoNuSeg | 2018 | Histopathology | Nuclei | Instance | 44 | https://monuseg.grand-challenge.org/ |
| GlaS | 2015 | Histopathology | Glands | Semantic/Instance | 85 | https://www.kaggle.com/datasets/sani84/glasmiccai2015-gland-segmentation |
| CoNIC | 2022 | Histopathology | Nuclei | Panoptic | 4,981 | https://conic-challenge.grand-challenge.org/Home/ |
| SegPC-2021 | 2021 | Microscopy | Plasma Cell | Instance | 490 | https://segpc-2021.grand-challenge.org/SegPC-2021/ |

*Continued on next page*

| Benchmark | Year | Modality | Segmentation Target | Segmentation Type | Scale | Link |
|---|---|---|---|---|---|---|
| HuBMAP-Hacking-the-Kidney | 2020 | Histopathology | Glomeruli | Instance | 20 | https://www.kaggle.com/competitions/hubmap-kidney-segmentation |
| HuBMAP+HPA-Hacking-the-Human-Body | 2022 | Histopathology | Functional Tissue Units | Instance | 880 | https://www.kaggle.com/competitions/hubmap-organ-segmentation |
| COSAS-Task1 | 2024 | Histopathology | Adenocarcinoma Regions | Semantic | 200 | https://cosas.grand-challenge.org/cosas/ |
| KPIs-2024 | 2024 | Histopathology | Glomeruli | Instance | $\geq 60$ | https://sites.google.com/view/kpis2024 |
| NeurIPS 2022 Cell Segmentation Competition Dataset | 2022 | Optical Microscopy | Cell | Instance | 1551 | https://zenodo.org/records/10719375 |
| MitoEM | 2021 | Electron Microscopy | Mitochondria | Instance | 1000 | https://mitoem.grand-challenge.org/ |
| MitoEM 2.0 | 2025 | Electron Microscopy | Mitochondria | Instance | ¿8000 | https://github.com/luckieucas/MitoEM2.0 |
| ISIC-Task1 | 2016<br>2017<br>2018 | Dermatoscope | Skin Lesion | Semantic | 900<br>2000<br>2700 | https://challenge.isic-archive.com/data/#2016<br>https://challenge.isic-archive.com/data/#2017<br>https://challenge.isic-archive.com/data/#2018 |
| CuRIOUS-SEG | 2022 | Ultrasound | Brain tumor and resection cavity | Semantic | 23 | https://curious2022.grand-challenge.org/ |
| Ultrasound Nerve Segmentation | 2016 | Ultrasound | Brachial Plexus | Semantic | 5635 | https://www.kaggle.com/competitions/ultrasound-nerve-segmentation |
| HC18 | 2018 | Ultrasound | Fetal head | Semantic | 999 | https://hc18.grand-challenge.org/ |
| CAMUS | 2019 | Ultrasound | Heart Structure | Semantic | 500 | https://www.creatis.insa-lyon.fr/Challenge/camus/ |
| TN-SCUI | 2020 | Ultrasound | Thyroid nodule | Semantic | 3644 | https://tn-scui2020.grand-challenge.org/ |
| FH-PS-AOP | 2023 | Ultrasound | Pubic symphysis & Fetal Head | Semantic | 4000 | https://ps-fh-aop-2023.grand-challenge.org/ |
| IUGC-Task2 | 2024 | Ultrasound | Pubic symphysis & Fetal Head | Semantic | 1000 | https://codalab.lisn.upsaclay.fr/competitions/18413 |
| UUSIC25 | 2025 | Ultrasound | Multi-organ segmentation | Semantic | 12002 | https://uusic2025.github.io/ |
| BKAI-IGH NeoPolyp | 2021 | Endoscopy | Polyp | Semantic | 1000 | https://www.kaggle.com/c/bkai-igh-neopolyp/ |
| GIANA (EndoVisSub) | 2017 | Endoscopy | Polyp | Semantic | 356 | https://endovissub2017-giana.grand-challenge.org/ |
| GIANA (EndoVisSub) | 2018 | Endoscopy | Polyp | Semantic | 356 | https://giana.grand-challenge.org/ |
| PolypGen | 2023 | Endoscopy | Polyp | Semantic | 8037 | https://endocv2021.grand-challenge.org/ |
| CHAOS | 2019 | MR/CT | Abdominal Organ | Semantic | 40 | https://chaos.grand-challenge.org/ |
| MSD | 2018 | MR/CT | Whole Body Organ/Lesion | Semantic | 1746 | http://medicaldecathlon.com/ |

*Continued on next page*

| Benchmark | Year | Modality | Segmentation Target | Segmentation Type | Scale | Link |
|---|---|---|---|---|---|---|
| MedSegBench | 2024 | Ultrasound, MRI, CT, X-ray, Endoscopy, Microscopy, Dermoscopy, Pathology | Multiple Tasks | Semantic | 53046 | https://doi.org/10.5281/ZENODO.13359660 |

Table 2: Summary of biomedical segmentation datasets.

| Dataset | Year | Modality | Segmentation Target | Segmentation Type | Scale | Link |
|---|---|---|---|---|---|---|
| MAMA-MIA | 2025 | MR | Primary Breast Tumor | Semantic | 1564 | https://www.ub.edu/mama-mia/ |
| Acute stroke annotated clinical MRI dataset | 2023 | MR | Brain Stroke | Semantic | 2888 | https://www.icpsr.umich.edu/web/ICPSR/studies/38464 |
| MOTUM | 2024 | MR | Brain Tumor | Semantic | 67 | https://dataverse.harvard.edu/dataset.xhtml?persistentId=doi:10.7910/DVN/KUUEWC |
| Longitudinal brain metastases dataset | 2025 | MR | Primary Breast Tumor | Semantic | 744 | https://doi.org/10.5281/zenodo.17253793 |
| CirrMRI600+ | 2025 | MR | Crrhotic Liver | Semantic | 628 | https://osf.io/cuk24/ |
| AbdomenCT-1K | 2021 | CT | Abdominal Organ | Semantic | 1000 | https://github.com/JunMa11/AbdomenCT-1K |
| RAOS | 2022 | CT | Abdominal Organ | Semantic | 413 | https://github.com/Luoxd1996/RAOS |
| AbdomenAtlas | 2023 | CT | Abdominal Organ | Semantic | 8448 | https://github.com/MrGiovanni/AbdomenAtlas |
| PLC-CECT | 2025 | CT | Liver Cancer Lesions | Semantic | 361 | https://www.scidb.cn/en/detail?dataSetId=9c0a7affa8154101b562a57e60ba9e96 |
| NLSTseg | 2025 | CT | Lung Tumor & Nodule | Semantic | 605 | https://zenodo.org/records/14838349 |
| CPAISD | 2025 | CT | Ischemic Core & penumbra | Semantic | 112 | https://zenodo.org/records/10892316 |
| COVID-QU-Ex | 2022 | X-ray | COVID-19 infection | Semantic | 33920 | https://www.kaggle.com/datasets/anasmohammedtahir/covidqu |
| CheXmask | 2024 | X-ray | Chest Anatomy | Semantic | 657566 | https://github.com/ngaggion/CheXmask-Database |
| FracAtlas | 2023 | X-ray | Bone Fracture | Instance | 4083 | https://github.com/XLR8-07/FracAtlas |
| DenPAR | 2025 | X-ray | Teeth | Semantic/Instance | 1000 | https://zenodo.org/records/13998619 |
| MTDDH | 2025 | X-ray | Pelvic Bones | Semantic | 1040 | https://www.scidb.cn/en/detail?dataSetId=c088644bd0b2406eb49830ad447c17fb |
| FIVES | 2022 | Fundus Photography | Retinal Vessel | Semantic | 800 | https://figshare.com/articles/figure/FIVES_A_Fundus_Image_Dataset_for_AI-based_Vessel_Segmentation/19688169 |
| PAPILA | 2022 | Fundus Photography | Optic Disc & Cup | Semantic | 488 | https://figshare.com/articles/dataset/PAPILA/14798004 |
| Chaksu | 2023 | Fundus Photography | Optic Disc & Cup | Semantic | 1345 | https://doi.org/10.6084/m9.figshare.20123135 |
| MAPLES-DR | 2024 | Fundus Photography | Diabetic Retinopathy Lesions & Anatomy | Semantic | 198 | https://doi.org/10.6084/m9.figshare.24328660 |
| PALM | 2024 | Fundus Photography | Pathologic Myopia Lesions & Optic Disc | Semantic | 1200 | https://doi.org/10.6084/m9.figshare.c.6224616.v1 |
| Leuven-Haifa | 2024 | Fundus Photography | Retinal Vessel | Semantic | 212 | https://doi.org/10.48804/Z7SHGO |

*Continued on next page*

| Dataset | Year | Modality | Segmentation Target | Segmentation Type | Scale | Link |
|---|---|---|---|---|---|---|
| Fundus-AVSeg | 2025 | Fundus Photography | Retinal Artery & Vein Vessel | Semantic | 100 | https://figshare.com/articles/dataset/Fundus-AVSeg/27938034 |
| PanNuke | 2020 | Histopathology | Nuclei | Panoptic | 7904 | https://github.com/Mr-TalhaIlyas/Prerpcessing-PanNuke-Nuclei-Instance-Segmentation-Dataset |
| NuInsSeg | 2024 | Histopathology | Nuclei | Instance | 665 | https://doi.org/10.5281/zenodo.10518968 |
| Nuclei-10Cancers | 2020 | Histopathology | Nuclei | Instance | 1356 | https://doi.org/10.6084/m9.figshare.12377135 |
| Data-Science-Bowl-2018 | 2019 | Optical Microscopy | Nuclei | Instance | 841 | https://bbbc.broadinstitute.org/BBBC038/ |
| LIVECell | 2021 | Optical Microscopy | Cells | Instance | 5239 | https://doi.org/10.6084/m9.figshare.14931555 |
| Kromp-Fluorescence-Nuclei | 2020 | Optical Microscopy | Nuclei | Instance | 79 | https://identifiers.org/biostudies:S-BSST265 |
| APACS23 | 2024 | Optical Microscopy | Cervical cells | Instance | 3565 | https://doi.org/10.17605/OSF.IO/CKA2F |
| PW-BALFC | 2025 | Optical Microscopy | BALF cells | Instance | 2105 | https://doi.org/10.5281/zenodo.14871206 |
| CellSAM | 2025 | Optical Microscopy | Cell | Instance | 10202 | https://vanvalenlab.github.io/cellSAM/API-key.html |
| TissueNet | 2022 | Optical Microscopy | Whole Cell & Nuclei | Instance | 3200 | https://deepcell.readthedocs.io/en/master/data-gallery/tissuenet.html |
| CREMI | 2016 | ssTEM | Neurites & Synapses | Semantic/Instance | 750 | https://service.tib.eu/ldmservice/dataset/cremi-dataset |
| SNEMI3D | 2013 | SEM | Neurites | Instance | 200 | https://zenodo.org/records/7142003 |
| UroCell | 2022 | FIB-SEM | Mitochondria & Lysosomes | Semantic | 1056 | https://github.com/MancaZerovnikMekuc/UroCell |
| PSFHS | 2024 | Ultrasound | Pubic symphysis & Fetal Head | Semantic | 1358 | https://zenodo.org/records/10969427 |
| ReMIND | 2024 | Ultrasound | Brain Tumor & Resection Cavity | Semantic | 114 | https://doi.org/10.7937/3RAG-D070 |
| High-Resolution Mouse Brain Tumor Ultrasound (Scientific Data) | 2025 | Ultrasound | Mouse brain tumor | Semantic | 1856 | https://doi.org/10.6084/m9.figshare.27237894 |
| BUS-UCLM | 2025 | Ultrasound | Breast lesion | Semantic | 683 | https://doi.org/10.17632/7fvgj4jsp7.3 |
| Spinal cord injury ultrasound dataset (Scientific Reports) | 2025 | Ultrasound | Spinal Cord Anatomy | Semantic | 10223 | http://github.com/HEPIUSLAB/ultrasound_spinal_cord_dataset |
| HyperKvasir (segmentation subset) | 2020 | Endoscopy | Polyp | Semantic | 1000 | https://doi.org/10.17605/OSF.IO/MH9SJ |
| Kvasir-SEG | 2020 | Endoscopy | Polyp | Semantic | 1000 | https://datasets.simula.no/kvasir-seg/ |
| ROBUST-MIS | 2019 | Endoscopy | Surgical Instruments | Instance | 10040 | https://www.synapse.org/Synapse:syn18779624 |
| PhaKIR | 2024 | Endoscopy | Surgical Instruments | Instance | 19435 | https://phakir.re-mic.de/ |

*Continued on next page*

| Dataset | Year | Modality | Segmentation Target | Segmentation Type | Scale | Link |
|---|---|---|---|---|---|---|
| Dresden Surgical Anatomy Dataset | 2023 | Endoscopy | Organs & Anatomical Structures | Semantic | 13195 | https://doi.org/10.6084/m9.figshare.21702600 |

### 2 Datasets for Foundation Model Training and Evaluation

The success of foundation models in medical image segmentation fundamentally depends on the scale, diversity, and quality of datasets used for pre-training and evaluation [1, 2]. While general-purpose models like SAM leverage natural image datasets such as SA-1B [3], their direct application to medical imaging is limited by the domain gap between natural and medical images—characterized by distinct visual properties, specialized anatomical structures, and varied imaging modalities [4, 5]. This has motivated the development of large-scale medical datasets specifically curated for training and evaluating foundation models, ranging from multi-modal collections spanning CT, MRI, and ultrasound to modality-specific datasets targeting particular clinical applications [6, 7, 8, 9]. In this section, we review the key datasets that underpin medical foundation model development, examining their characteristics, coverage, and role in enabling generalizable segmentation capabilities.

#### 2.1 2D Medical Image Datasets

Among the diverse medical imaging resources, two-dimensional (2D) datasets constitute the majority and remain pivotal for foundation model development. Representative examples include CT- and MRI-based datasets such as RSNA Intracranial Hemorrhage Detection [10] ($\approx$ 900k brain CT slices) and RadImageNet [11] (multimodal CT–MRI coverage), which provide large-scale, anatomically diverse data for classification and detection. X-ray collections such as CheXpert [12] (224k chest radiographs), NIH ChestX-ray14 [13] (112k), and CheXmask [14] (670k lung masks) further enable multi-label and segmentation training, while ophthalmic datasets OCT2017 [15] and EyePACS [16], together with dermatology benchmarks like the ISIC series [17], expand large-scale resources for retinal and dermoscopic disease analysis. Collectively, these 2D datasets illustrate both the breadth and fragmentation of current medical imaging resources: large-scale collections exist for a few clinically prioritized domains, yet many anatomical regions remain underrepresented. Segmentation serves as the most critical task supported by these datasets, providing pixel-level delineation of organs and lesions essential for downstream applications such as treatment planning and disease monitoring. Despite uneven coverage, these datasets form indispensable testbeds for developing and benchmarking foundation models, enabling evaluation of model generalization across modalities, tasks, and clinical settings.

#### 2.2 3D Medical Image Datasets

In contrast to the abundance of 2D datasets, three-dimensional (3D) medical imaging resources are fewer but hold greater clinical value, with CT and MRI providing indispensable volumetric context for diagnosis and treatment planning. Key examples include TotalSegmentator [18] (1,000+ annotated CT volumes across 104 structures) and CT-RATE [19] (25,000+ chest CT scans), which enable large-scale multi-organ segmentation and drive volumetric foundation model development. Nonetheless, most 3D datasets remain relatively small and organ-specific, as seen in BraTS [20] for brain tumor and AMOS [21] for abdominal segmentation, with growth constrained by the high cost of acquisition, annotation, and computational demands. Despite these limitations, 3D datasets are indispensable for advancing medical foundation models, as volumetric segmentation offers richer spatial understanding than 2D data and provides essential testbeds for assessing generalization across complex, multi-organ, and multi-modality tasks.

#### 2.3 Video Medical Image Datasets

Compared with 2D and 3D imaging, video datasets are fewer but uniquely enable temporal modeling for dynamic clinical segmentation tasks, particularly in surgical and endoscopic domains where frame-level tool and organ delineation is critical. The EndoVis challenge series provide laparoscopic videos with detailed instrument segmentation and tracking labels, establishing the benchmark for computer-assisted surgery, while Cholec80 [22] extends this direction with 80 annotated cholecystectomy videos for workflow recognition and segmentation. In gastrointestinal imaging, Kvasir-SEG [23], Hyper-Kvasir [24], and GIANA datasets offer large-scale colonoscopy recordings with polyp masks, whereas ophthalmic and robotic datasets such as Cataract-101/Cataract-1K [25] and SARAS-MESAD [26] provide densely annotated videos for surgical phase recognition and multi-camera tool segmentation. Together, these datasets establish segmentation as the central task for video-based benchmarks, enabling the development of foundation models that can operate reliably in real-time, temporally coherent clinical environments.

### 3 Adaptation of Foundation Models for Biomedical Image Segmentation

#### 3.1 Adaptation of Foundation Models for Biomedical Image Segmentation

Vision foundation models such as the Segment Anything Model (SAM) have revolutionized image segmentation by introducing a promptable interface and enabling powerful zero-shot generalization across diverse visual domains [27, 28]. Yet, when directly applied to medical imaging, prompting alone often proves inadequate. Medical images are characterized by low contrast and subtle boundaries [29, 30, 31], diverse modalities with distinct signal properties [32, 33], and anatomically complex structures [34, 35] that differ fundamentally from natural scenes. These factors highlight a critical gap between promptable generalization and clinical reliability, motivating the development of comprehensive adaptation strategies that extend beyond prompting. Recent advances include fine-tuning frameworks that recalibrate model parameters for domain-specific representations [29, 36], architectural refinements that enhance multi-scale and volumetric reasoning [37], and optimization schemes tailored to medical imaging constraints [35]. Together, these efforts mark a transition from generic promptable segmentation toward clinically adaptive foundation models capable of meeting the stringent accuracy, robustness, and interpretability requirements of medical practice. Building on this motivation, the following sections review the principal directions for adapting and extending vision foundation models in medical image segmentation.

##### 3.1.1 Fine-tuning

Despite SAM's broad generalization ability, extensive benchmarks reveal that its zero-shot transfer to medical imaging remains limited. While it performs competitively on large and clearly defined structures with sufficient prompting [38, 39], performance degrades sharply for low-contrast organs, tumors, and volumetric data [40, 41, 4]. These deficits highlight the need for systematic adaptation through fine-tuning, which has emerged as the dominant strategy for aligning general-purpose VFMs with medical tasks.

Early studies demonstrate that full fine-tuning of SAM yields the strongest performance across modalities. MedSAM [29] and its successors, including SAM-Med2D [42], SAM-Med3D [43], AutoSAM [44], Biomedical SAM 2 [45], and MedSAM2 [46], consistently demonstrate that large-scale retraining closes the domain gap and enables stable 2D, 3D, and video segmentation. However, such approaches demand enormous computational and data resources, limiting their practicality outside centralized research infrastructures.

To mitigate these costs, parameter-efficient fine-tuning (PEFT) strategies have become the preferred alternative. LoRA-based variants such as SAMed [47], PTSAM [48], BLO-SAM [49], FLAP-SAM [50], Conv-LoRA [51], and their task-specific extensions [52, 53, 54, 55, 56, 57] reveal that selective adaptation of encoders and decoders can recover much of the accuracy of full retraining while preserving pretrained generality. These methods mark a pragmatic balance between scalability and performance, enabling rapid specialization to new organs or modalities. Building on this efficiency-oriented perspective, recent work moves beyond lightweight parameter tuning to explore deeper architectural coupling, aiming to endow SAM with richer representations and stronger domain reasoning. Beyond LoRA, hybrid architectures fuse SAM with complementary learners to enhance representation richness and reduce reliance on prompts [58, 59, 60, 61, 62, 63, 64, 65, 66]. Knowledge distillation and uncertainty modeling are increasingly adopted to transfer SAM's representation power to lightweight or probabilistic variants, promoting efficiency and interpretability.

Building upon parameter-efficient strategies, adapter-based methods introduce lightweight modules that enable flexible domain adaptation while preserving the core SAM architecture. Rather than updating all parameters, these approaches insert learnable components—such as prompt generators, attention bridges, or domain-aware adapters—to endow SAM with task-specific reasoning at low computational cost. A major focus lies in automating the prompt interface itself. Works including AutoSAM [44], TongueSAM [67], Med-PerSAM [68], SAMUS [69], EchoONE [70], and MedLSAM [71] replace manual interactions with automatic prompt generation or localization. Recent works further integrate external knowledge sources such as CLIP or foundation models, as seen in SaLIP [72], SAM-Path [73], and MCP-MedSAM [74], which synthesize visual and textual representations to improve segmentation consistency. Others emphasize self-prompting and task embedding to achieve prompt-free inference [75, 76, 77]. Collectively, these efforts demonstrate a clear trajectory from user-dependent interaction to autonomous, context-aware prompting—signifying a shift toward self-reasoning segmentation systems that generalize across modalities.

Recent advances enhance SAM's reasoning and contextual understanding by integrating memory and attention mechanisms that capture spatial, temporal, and cross-modal dependencies critical for medical data. MedSAM-2 [78] and SAMed-2 [79] exemplify this trend through self-sorting memory banks and temporal adapters that stabilize multi-task learning and mitigate catastrophic forgetting, while memory-augmented designs such as Memorizing SAM [80], MemSAM [81], and SAM4EM [82] preserve spatial-temporal coherence in volumetric and video segmentation. Beyond explicit memory, attention-centric architectures—including ProMISe [83], Med-SAM-Adapter [84], and BiaSAM [85]—refine token interactions via lightweight or frequency-aware attention, while BUSSAM [86] and CC-SAM [87] employ multi-branch and variational attention to enhance feature localization in ultrasound and radiological imaging. Further extending this idea, cross-modality adapters for brain MRI [88], Hermes [89], and CD-SAM [90] unify attention across modalities, tasks, and 2.5D spatial contexts, achieving more consistent segmentation under complex anatomical variability. These designs illustrate a broader shift toward memory- and attention-augmented SAM variants that reason across space, time, and modality—paving the way for adaptive, context-aware medical segmentation.

Building on these advances, domain-specific and training-framework adapters further extend SAM's adaptability by embedding medical priors and refining its learning dynamics for real-world deployment. Domain-oriented methods such as EMedSAM [91] and LeSAM [92] introduce adaptive modules that encode clinical priors and lesion morphology, while UA-SAM [93] models annotation ambiguity through uncertainty-aware adaptation. SAMIHS [94] enhances low-contrast CT segmentation via boundary-sensitive adapters, and Trans-SAM [95] injects convolutional inductive biases for efficient cross-modality transfer. Hierarchical extensions such as SAM2-Adapter [96] and contrastive text-vision fusion frameworks like [97] further enrich representational depth and robustness under noisy conditions. Complementing these domain-focused strategies, training-framework adapters—such as meta-learning and federated paradigms in SSM-SAM [98] and FedFMS [99]—enable fast adaptation and privacy-preserving collaboration, while unified and semi-supervised systems including SAMME [100], ASLseg [101], PP-SAM [102], and $\mu$-SAM [103] reduce annotation demands via pseudo-labeling and iterative refinement. Online and continual learning frameworks such as [104] and AuxOL [105] extend SAM's capacity to integrate real-time feedback, positioning it as a continuously learning, domain-aware foundation model capable of adaptive performance across modalities, institutions, and evolving clinical settings.

##### 3.1.2 Architecture Modification

The adaptation of SAM for medical image segmentation has driven extensive architectural innovation, reflecting a broader movement from 2D, prompt-dependent design toward 3D, anatomy-aware, and contextually grounded segmentation. These efforts range from radical reconstructions of SAM's architecture to lightweight refinements that enhance efficiency and structural understanding. The central motivation lies in addressing SAM's inherent limitations in volumetric reasoning, cross-slice consistency, and domain-specific feature extraction—critical challenges in medical imaging. By expanding SAM's representational depth and integrating medical priors, these architectural advances not only improve segmentation accuracy but also redefine how foundation models can generalize across modalities and clinical tasks.

Recent works diverge along three primary directions: core architectural redesign, hybrid integration and adapter-based modification. Recent advances showcase diverse yet convergent strategies for achieving this goal. Full 3D reconstructions such as SAM-Med3D [106], SAM-Med3D-MoE [107], SAM3D [108], SAM3X [109], CT-SAM [110], SegVol [111], and CT-SAM3D [112] replace SAM's slice-based reasoning with fully volumetric or multi-axis encoders, substantially improving anatomical consistency but often at the expense of higher compute and limited interactivity. Efficiency-oriented designs like FastSAM3D [113], VSS-SAM [114], and nnInteractive [115] counterbalance this by compressing attention layers or integrating lightweight CNN or Mamba branches, achieving real-time or interactive 3D inference without major accuracy loss. A parallel trend focuses on embedding anatomical priors and multimodal reasoning—exemplified by cineCMR-SAM [116], ASAM [117], BrainSegDMIF [118], and SyncSAM [119]—which integrate temporal, cross-modality, or Fourier-domain cues for improved generalization. Meanwhile, refined architectures—such as DeSAM [120], SIT-SAM [121], H-SAM [122], and UN-SAM [123]—retain SAM's encoder but introduce hierarchical decoding, self-prompting, or semantic transformers to enhance zero-shot and prompt-free segmentation. Extending these ideas, recent SAM2-based models like FATE-SAM [124], SLM-SAM2 [125], SurgSAM2 [126], and SAM2Med3D [127] incorporate explicit memory and temporal reasoning for video and 3D workflows, achieving improved spatial-temporal consistency. Finally, foundation-level extensions such as VesselFM [128], VISTA3D [129], and SegmentAnyBone [130] push toward universal 3D segmentation backbones that unify interactive and automatic workflows.

An emerging research direction focuses on hybrid integration frameworks that fuse SAM's transformer-based representation with complementary architectures to improve adaptability, efficiency, and multimodal reasoning across medical domains. These approaches combine SAM's global perception with localized feature extraction to recover structural precision often lost in pure ViT models. Early efforts such as SAMDA [131], WSI-SAM [132], and MoME [133] highlight the benefits of coupling SAM with CNNs or modality experts for histopathology and MRI, achieving superior cross-domain generalization. Similar hybrid designs—ASPS [134], MBA-Net [135], and MedSAM-CA [136]—enhance boundary fidelity through multi-scale fusion and uncertainty-guided refinement, while models like SpineFM [137], SAM-FNet [138], and SAM-Swin [139] integrate SAM-derived priors for lesion and bone segmentation, demonstrating improved interpretability and robustness. More recent advances extend hybridization to SAM2, as seen in SAMba-UNet [140] and Path-SAM2 [141], which merge Mamba or UNI encoders for stronger long-range dependency modeling. Meanwhile, SAM-MIL [142], Slide-SAM [143], and SegAnyPET [144] adapt SAM's prompt mechanisms to weakly supervised, volumetric, or low-signal modalities, underscoring its versatility beyond full supervision. Rep-MedSAM [145] takes a direct approach by completely replacing the original image encoder with RepViT, specifically designed for mobile-friendly deployment scenarios. These hybrid systems reveal a transition from monolithic segmentation models to compositional medical foundation architectures—where SAM acts as a unifying representational core enhanced by domain-aware, modality-specific, and efficiency-driven modules to meet the precision and scalability demands of real-world medical imaging.

**Adapter-Based Modification.** A parallel trend focuses on adapter-based modification, where lightweight modules are inserted into SAM's architecture to enable efficient domain adaptation without full retraining. These adapters aim to preserve SAM's general visual priors while injecting modality- and task-specific information, striking a balance between flexibility and parameter efficiency. Early efforts such as MedSAM-Adapter (Med-SA) [146] and DASAM [147] highlight the benefits of space-depth adaptation and simplified prompt tuning, achieving substantial performance gains with minimal parameter cost. SPA [148] and I-MedSAM [149] further refine this paradigm through convolutional and frequency-domain adapters that enhance multi-scale and uncertainty-aware reasoning, illustrating how structural priors can be integrated without compromising generality. Recent works like M-SAM [150] and MA-SAM [5] expand adapter tuning into the 3D and temporal domains, demonstrating that partial adaptation can rival or surpass fully fine-tuned models in volumetric tasks. SAM2-UNet [151], Tri-Plane Mamba [152], and FreqSAM2-UNet [153] further exemplify this direction by embedding adapters into hierarchical or frequency-aware backbones to enhance context modeling across scales. Complementarily, MASG-SAM [154] integrates hierarchical attention and boundary-aware adapters to improve few-shot segmentation, underscoring the growing emphasis on structural efficiency and generalization. Taken together, these designs signal a broader shift from architecture-heavy fine-tuning toward modular adaptability, demonstrating that minimal yet strategically placed adapters can endow SAM with cross-modality flexibility, reduced training cost, and competitive accuracy in medical segmentation.

### 4 Architectural Details of U-Net Variants

#### 4.1 Skip Connection Variants

As shown in Fig. 1(a), the evolution of U-Net architectures reflects a progressive redefinition of skip connections from simple feature transfer mechanisms to structured modules for semantic alignment and multi-scale fusion. In the original U-Net, skip connections directly concatenate encoder and decoder features at matching resolutions, effectively preserving spatial detail but offering limited capability to reconcile the semantic disparity between low-level and high-level representations.

Subsequent variants enhance skip connections by increasing their structural depth and connectivity. U-Net++ [155] replaces direct connections with nested and densely connected skip pathways, introducing intermediate convolutional transformations that gradually bridge semantic gaps. This design enables deep supervision and produces segmentation outputs at multiple semantic levels, improving optimization stability and feature coherence across the network.

Extending this idea, U-Net 3+ [156] adopts full-scale skip connections, allowing each decoder stage to aggregate features from all encoder levels and preceding decoder layers. This dense inter-scale integration ensures that fine spatial details and global context jointly inform decoding, while channel compression maintains computational efficiency.

Other approaches focus on enriching the representational capacity within skip pathways themselves. SIU-Net [157] embeds inception-style multi-scale convolutions into skip connections, enabling scale-adaptive feature extraction prior to fusion. This design is particularly effective for segmenting objects with large size variability. Multi-stage architectures further extend the functional role of skip connections. DoubleU-Net [158] introduces a cascaded refinement strategy in which the second U-Net leverages both the initial segmentation and intermediate features from the first network. Similarly, W-Net [159] employs dual U-Nets to jointly enforce segmentation and reconstruction consistency, using skip connections as implicit structural regularizers.

Skip connections have also been augmented with explicit contextual modeling. Context-aware U-Net [160] integrates neighboring-region context modules into skip pathways, enlarging the effective receptive field and improving discrimination in locally ambiguous regions without increasing network depth.

#### 4.2 Encoder Variants

As summarized in Fig. 1(b), architectural advances in U-Net encoders primarily focus on improving feature extraction depth, enhancing gradient flow, and strengthening multi-scale context modeling. While the original U-Net encoder relies on stacked convolution–pooling blocks, this design limits representational capacity as network depth increases and may lead to information loss in fine structures.

A major line of development introduces residual and dense connectivity to stabilize optimization and promote feature reuse. ResUNet [161] replaces standard convolutional blocks with residual units, enabling deeper encoders by alleviating degradation and vanishing gradient issues. This residual formulation preserves low-level features across layers and improves robustness in noisy medical images. ResUNet++ [162] further enhances this design by integrating Squeeze-and-Excitation blocks for channel-wise attention and Atrous Spatial Pyramid Pooling for multi-scale context aggregation, allowing the encoder to adaptively emphasize informative features while capturing scale variability. In contrast, Dense U-Net [163] adopts DenseNet-style dense blocks, concatenating features from all preceding layers within each encoder stage. This dense connectivity maximizes feature reuse and strengthens gradient propagation, proving effective for preserving thin and curvilinear structures.

Beyond connectivity patterns, several variants redesign the internal structure of encoder blocks to achieve hierarchical multi-scale representation. U2-Net [164] introduces Residual U-blocks, embedding mini U-Net structures within each encoder stage to capture both local and global context without aggressive downsampling. This nested design maintains spatial resolution while increasing effective depth and enables deep supervision through side outputs. Similarly, DRU-Net [165] employs dilated residual units to expand the receptive field without reducing feature map resolution, a property particularly beneficial for high-precision tasks such as OCT layer segmentation.

Multi-scale feature extraction is further emphasized through parallel convolutional designs. MSU-Net [166] incorporates multi-receptive-field blocks directly into the encoder, enabling scale-aware feature learning at each stage. Inception U-Net [167] follows a similar strategy by replacing standard convolutions with inception modules, while Dense-Inception U-Net [168] combines inception-style multi-scale processing with dense connectivity to jointly enhance contextual diversity and feature reuse.

#### 4.3 Decoder Variants

As illustrated in Fig. 1(c), architectural refinements in U-Net decoders and bottlenecks primarily aim to improve feature reconstruction, boundary precision, and semantic consistency during upsampling. While early U-Net designs rely on symmetric decoder paths with simple upsampling and concatenation, such strategies often struggle to recover fine structures and sharp boundaries, especially when encoder representations become increasingly abstract.

A prominent direction enhances feature selection and refinement within the decoder. RIC-Unet [169] introduces Deconvolution–Channel (DC) blocks that integrate channel attention into the upsampling process. By selectively reweighting feature channels through Channel Attention Blocks, the decoder emphasizes discriminative responses while suppressing noise, leading to more accurate boundary reconstruction in challenging settings such as histopathological nuclei segmentation. Similarly, MA-Net [170] incorporates Multi-scale Fusion Attention Blocks that jointly attend to low-level spatial details and high-level semantic features. This attention-guided fusion improves scale robustness and enables more precise delineation of structures with heterogeneous sizes.

Another line of work expands the functional role of the decoder through multi-task and auxiliary supervision. PSI-Net [171] adopts a multi-branch decoder that simultaneously predicts segmentation masks, contour maps, and distance transforms. This design explicitly encodes boundary and geometric priors into the reconstruction process, improving instance separation and contour smoothness by allowing complementary tasks to regularize one another. In volumetric segmentation, SegResNet [172] integrates a variational autoencoder branch at the bottleneck, introducing an auxiliary reconstruction objective that constrains latent representations. This bottleneck regularization improves generalization, particularly in data-limited 3D medical imaging scenarios, by encouraging the encoder–decoder pathway to retain anatomically meaningful information.

In addition to specialized decoder designs, many architectures maintain encoder–decoder symmetry by extending enhanced encoder blocks into the decoding path. Models such as ResUNet [161], Dense U-Net [163], and Inception U-Net [167] apply residual, dense, or inception-style modules consistently across both stages, ensuring coherent feature transformations during both abstraction and reconstruction.

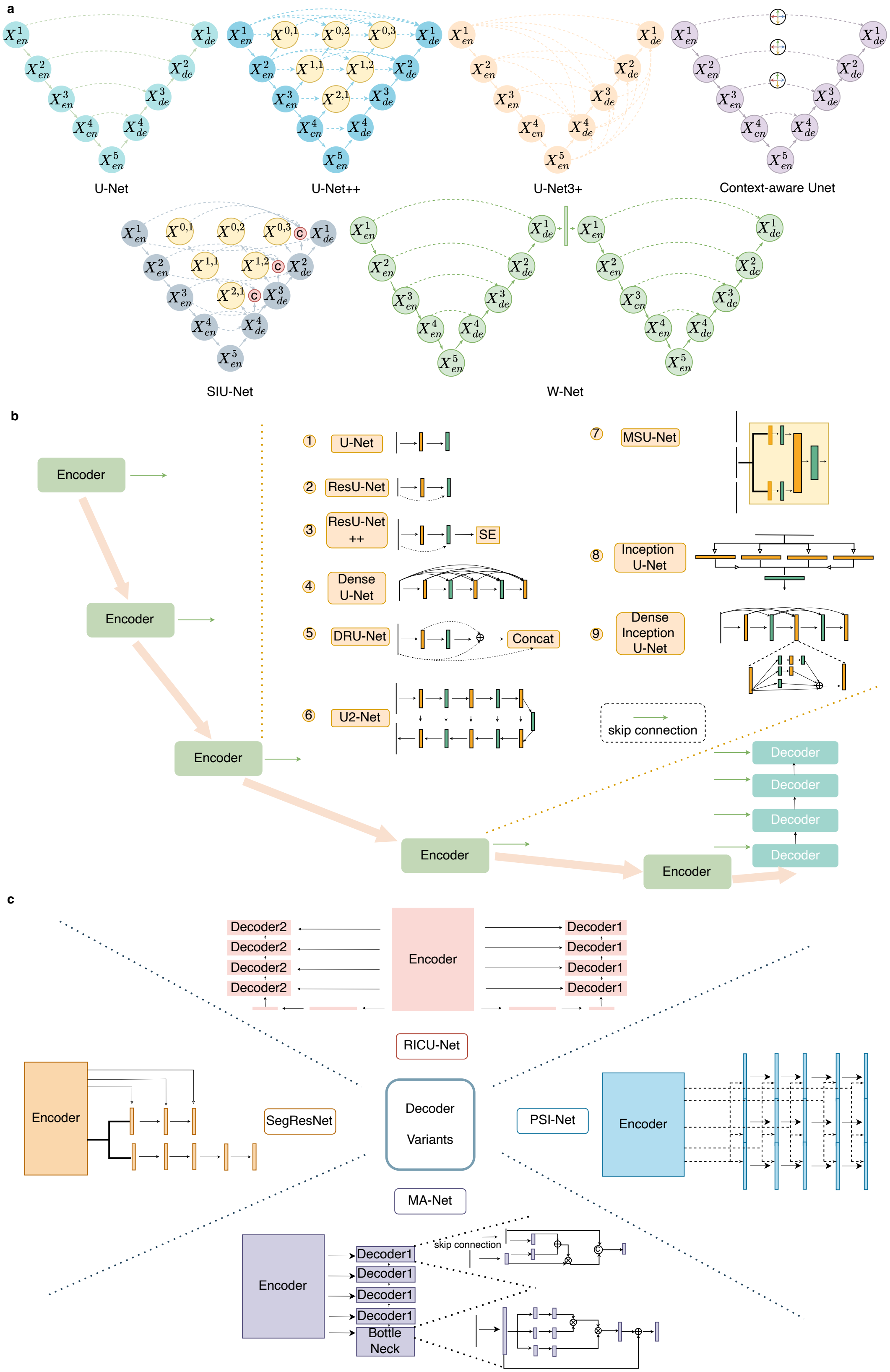


Figure 1: Architectural evolution of representative U-Net variants.

#### 4.4 Hybrid CNN-Attention models

As illustrated in Fig. 2(a), hybrid CNN–attention architectures represent an intermediate stage between purely convolutional U-Net variants and fully Transformer-based models. These approaches aim to augment the strong local feature extraction and inductive bias of CNNs with attention mechanisms that selectively emphasize relevant structures or incorporate broader contextual information, while avoiding the computational cost and data requirements of full Transformers.

Early hybrids integrate lightweight attention modules directly into U-Net-style architectures. Attention U-Net [173] introduces attention gates along skip connections to suppress irrelevant activations and highlight salient anatomical regions, improving sensitivity to small or low-contrast structures. This idea is further extended in Residual Attention U-Net [174], which combines attention gating with residual blocks to stabilize deeper networks and sharpen boundary transitions. Attention U-Net++ [175] embeds attention into the nested skip pathways of UNet++, narrowing semantic gaps and yielding more spatially coherent predictions. Domain-adapted designs such as ASCU-Net [176] tailor attention to lesion-specific cues, while USE-Net [177] employs Squeeze-and-Excitation–style channel attention to recalibrate feature responses and improve robustness across datasets.

Building on these CNN-centric attention designs, later hybrids seek to combine convolutional encoders with Transformer-style self-attention for global context modeling. A common strategy inserts a Transformer at the bottleneck of a U-shaped network. TransUNet [178] couples a CNN encoder with a ViT, fusing tokenized global representations with high-resolution decoder features. In volumetric settings, TransBTS [179] and Swin-UNETR [180] replace ViT with more efficient or hierarchical Transformers, enabling long-range dependency modeling while preserving spatial structure.

Another design axis concerns where and how attention is fused. Bottleneck-centric approaches such as TransAttUnet [181] and AFTer-UNet [182] inject axial or multi-level attention between encoder and decoder, whereas U-Transformer [183] and UCATR [184] extend self- and cross-attention to skip connections and deeper pathways. Parallel fusion strategies also emerge: TransFuse [185] processes images through CNN and Transformer branches simultaneously and merges them via BiFusion, while MCTrans [186] performs inter- and intra-scale attention over multi-scale tokens. Related designs such as DC-Net [187], UTNet [188], nnFormer [189], CoTr [190], and Mixed Transformer U-Net [191] explore alternative arrangements of convolution, attention, and skip interactions to balance efficiency and global reasoning.

Finally, several task-driven hybrids emphasize robust fusion under limited data or multi-modal inputs. Examples include Swin-UNet3D [192], X-Net [193], and multi-attention designs such as DATTNet [194], DA-TransUNet [195], and MedFuseNet [196], which combine spatial, channel, and Transformer-based attention to stabilize learning and improve generalization.

#### 4.5 Transformer-Only segmenters

As shown in Fig. 2(b), a parallel line of U-Net–inspired architectures abandons convolutional backbones almost entirely, relying instead on Transformer-based self-attention to model long-range dependencies and global context. These models aim to minimize convolutional inductive biases and enable more flexible representation learning, particularly for complex anatomical structures with large spatial variability.

Early Transformer-only designs retain the U-shaped paradigm while replacing convolutional blocks with hierarchical attention modules. Swin-Unet [197] employs a pure Swin-Transformer encoder–decoder, using shifted-window self-attention to balance global context modeling with computational efficiency. This fully Transformer-based formulation demonstrates strong performance across a wide range of organ and lesion segmentation tasks, indicating that hierarchical attention can effectively substitute for convolutional feature extraction. DS-TransUNet [198] extends this idea by introducing dual Swin-Transformer streams that separately encode local and global representations, enhancing contextual diversity and improving robustness to scale variation.

Beyond windowed attention, newer architectures explore alternative token interaction mechanisms to further improve efficiency and adaptability. D-Former [199] introduces dynamic token propagation, allowing information to be selectively routed across layers based on content relevance rather than fixed spatial neighborhoods. This dynamic formulation enhances long-range dependency modeling while reducing redundant computation. More recent designs such as SMAFormer [200] further enrich Transformer backbones by coupling pixel-level, channel-level, and spatial attention within a unified architecture, enabling fine-grained control over feature interactions at multiple representational levels.

### 5 Evaluation Protocols

Segmentation papers often report strong headline numbers while leaving evaluation details underspecified. In biomedical imaging, this is risky: small geometric deviations can change downstream measurements, and domain shift across scanners, sites, and annotators is frequently the dominant failure mode. We therefore recommend structuring evaluation along three orthogonal axes: *granularity* (semantic, instance, panoptic), *geometry* (overlap vs. boundary/surface), and *operational cost* (interaction burden, time, and robustness across sites). Below we summarize a practical metric set, define each quantity explicitly, and outline task-aware reporting protocols rather than prescribing a single universal score.

#### 5.1 Traditional Segmentation Metrics by Granularity

##### 5.1.1 Semantic Segmentation

Semantic segmentation operates at the region or class level and remains the workhorse setting for organ, lesion, and tissue compartment delineation. In biomedical imaging, it is typically the first step from which volumetric, morphometric, or radiotherapy planning quantities are derived, so both overlap and boundary behavior matter.

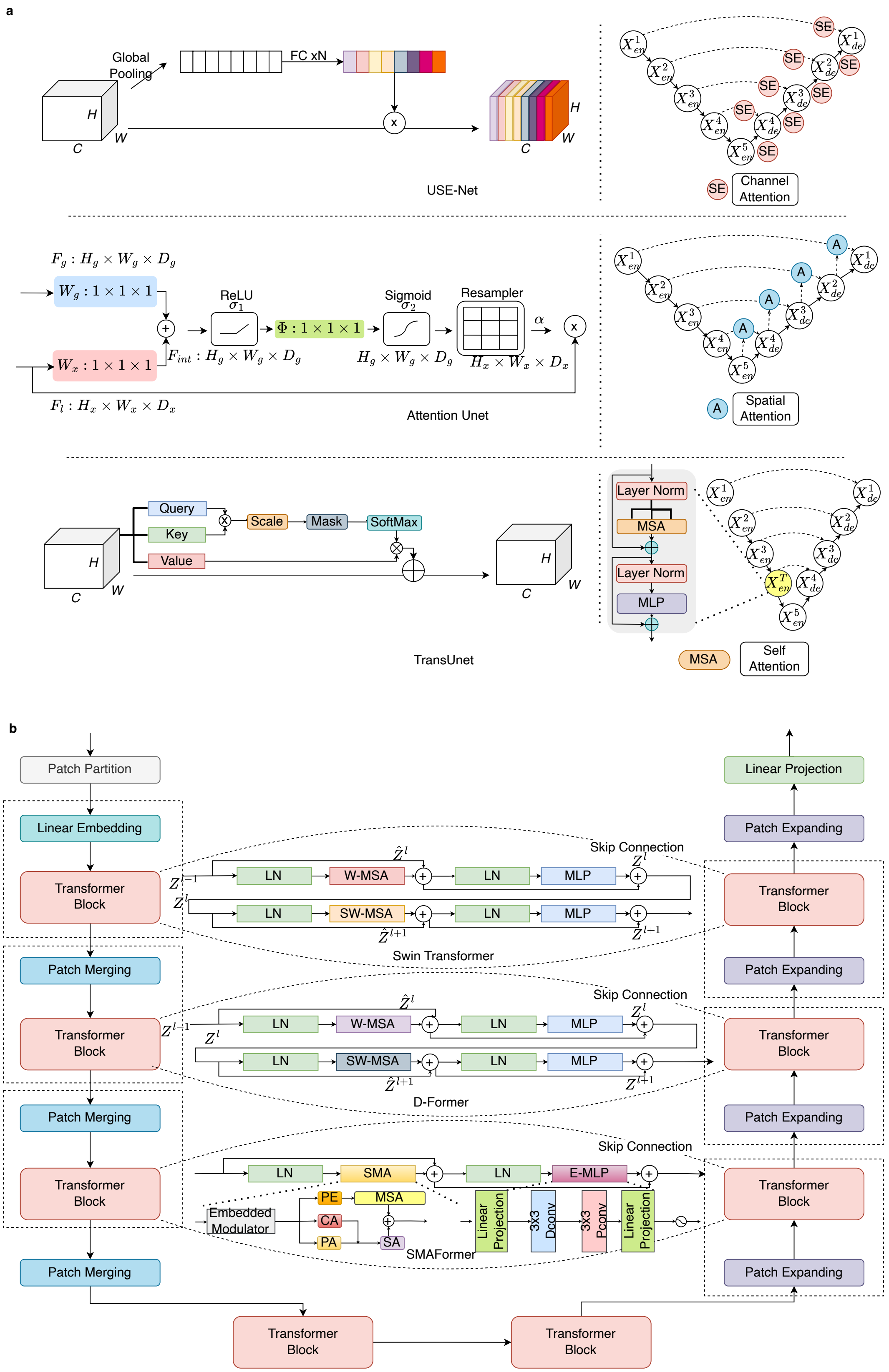


Figure 2: The evolution from CNN-based attention to Transformer-based foundational models.

Let $\Omega$ be the image domain. Let $Y \subseteq \Omega$ denote the ground-truth foreground set (or a class-specific set) and $\hat{Y} \subseteq \Omega$ the prediction. Define pixel-wise counts in the usual way:

$$TP = \left|\hat{Y} \cap Y\right|, \quad FP = \left|\hat{Y} \setminus Y\right|, \quad FN = \left|Y \setminus \hat{Y}\right|, \quad TN = \left|\Omega \setminus \left(\hat{Y} \cup Y\right)\right|.$$

**Overlap based core metrics.** The most commonly reported region metrics are the Dice similarity coefficient (Dice or DSC)[201] and the intersection over union (IoU)[201]:

$$\text{Dice}(\hat{Y}, Y) = \frac{2TP}{2TP + FP + FN}, \qquad \text{IoU}(\hat{Y}, Y) = \frac{TP}{TP + FP + FN}.$$

These capture how well foreground regions overlap but do not distinguish where errors occur or how they affect small structures.

**Multi class summaries under imbalance.** For multi class semantic segmentation, it is important to report both per class scores and class balanced summaries, especially when rare but clinically important classes are present. Let $\{Y_c\}_{c=1}^{C}$ and $\{\hat{Y}_c\}_{c=1}^{C}$ be class specific sets. The mean IoU is

$$\text{mIoU} = \frac{1}{C} \sum_{c=1}^{C} \text{IoU}(\hat{Y}_c, Y_c).$$

Class wise $F1_c$ and its macro average further summarize performance across classes:

$$\text{F1}_c = \frac{2 \, \text{Precision}_c \, \text{Recall}_c}{\text{Precision}_c + \text{Recall}_c}, \quad \text{MacroF1} = \frac{1}{C} \sum_{c=1}^{C} \text{F1}_c,$$

where for each class $c$

$$\text{Precision}_c = \frac{TP_c}{TP_c + FP_c}, \quad \text{Recall}_c = \frac{TP_c}{TP_c + FN_c}, \quad \text{Specificity}_c = \frac{TN_c}{TN_c + FP_c}, \quad \text{Accuracy}_c = \frac{TP_c + TN_c}{TP_c + TN_c + FP_c + FN_c}.$$

In practice, reporting both macro and micro summaries helps separate overall pixel dominance from performance on minority classes.

**Boundary and surface distances for clinically sensitive tasks.** In many biomedical applications, especially radiotherapy planning and surgical guidance, small boundary deviations can have larger impact than modest changes in volumetric overlap. For this reason, overlap metrics are often complemented by boundary and surface distance measures. Let $S_Y$ and $S_{\hat{Y}}$ denote the boundary (in 2D) or surface (in 3D) point sets of $Y$ and $\hat{Y}$, respectively. The point to surface distance is

$$d(x, S) = \min_{y \in S} \|x - y\|_2 \quad \text{for point } x \text{ and surface } S.$$

Using this definition, the Hausdorff distance (HD)[202] and average symmetric surface distance (ASSD)[202] are

$$\text{HD}(S_{\hat{Y}}, S_Y) = \max \left\{ \sup_{x \in S_{\hat{Y}}} d(x, S_Y), \; \sup_{y \in S_Y} d(y, S_{\hat{Y}}) \right\},$$

$$\text{ASSD}(S_{\hat{Y}}, S_Y) = \frac{1}{|S_{\hat{Y}}| + |S_Y|} \left( \sum_{x \in S_{\hat{Y}}} d(x, S_Y) + \sum_{y \in S_Y} d(y, S_{\hat{Y}}) \right).$$

In clinical studies, a robust variant such as the 95th percentile Hausdorff distance (HD95) is frequently preferred, obtained by replacing each sup with the 95th percentile of directed distances.

**Tolerance aware boundary agreement.** For tasks where there is an explicit notion of acceptable contouring error, tolerance based metrics are more interpretable. The surface Dice at tolerance $\tau$[203] measures the fraction of surface points that lie within $\tau$ of the opposite surface:

$$\text{SDice}_\tau = \frac{\left|\{x \in S_{\hat{Y}} : d(x, S_Y) \leq \tau\}\right| + \left|\{y \in S_Y : d(y, S_{\hat{Y}}) \leq \tau\}\right|}{|S_{\hat{Y}}| + |S_Y|}.$$

Here $\tau$ is chosen according to the spatial tolerance of the target application (for example a few millimeters in radiotherapy), allowing direct comparison to clinically acceptable deviation thresholds rather than abstract geometric error alone.

**Recommended protocol for semantic biomedical segmentation.** Rather than reporting a single headline Dice, we recommend a small, task-aware bundle of metrics:

- For organ and large-structure segmentation, report per-structure Dice and HD95 at the *case level*, together with summary statistics (mean, standard deviation, and quartiles) across cases.
- For radiotherapy and surgical planning, always complement region overlap (Dice or IoU) with boundary-aware metrics such as HD95 and $\text{SDice}_\tau$, with $\tau$ chosen to match clinically accepted contouring tolerances (for example a few millimeters for organs-at-risk).
- For class-imbalanced problems with rare but safety critical classes, report both micro and macro averages, and explicitly list per-class scores for the rare targets rather than hiding them in a global mean.
- When comparing methods, avoid reporting only pooled pixel-level summaries; instead, show distributions of case-level scores (for example box plots of Dice and HD95), which make robustness and failure rates visible.

These elements constitute a minimal semantic segmentation protocol that balances volumetric accuracy with boundary fidelity and class-level safety.

##### 5.1.2 Instance Segmentation

Instance segmentation is often the clinically meaningful granularity in biomedical imaging, because downstream use frequently depends on counting, sizing, and individually characterizing discrete structures (for example nuclei, glands, lesions, cells, micro-anatomical compartments). Evaluation therefore needs to separate (i) whether instances are found at all, (ii) whether one-to-one correspondence is correct (splits and merges), and (iii) whether the delineation is accurate once matched.

**Matching rule (object correspondence).** Most object-level metrics begin with a one-to-one matching between predicted instances $\{\hat{M}_i\}$ and ground-truth instances $\{M_j\}$, typically using an IoU threshold and a maximal matching (often via greedy IoU or Hungarian assignment):

$$\mathrm{IoU}(A,B) = \frac{|A \cap B|}{|A \cup B|}.$$

Let $\mathcal{T}$ be the set of matched pairs, and define object-level $TP, FP, FN$ after matching.

**Object detection quality (did we find the right instances?).** A simple and widely used summary is object-level precision/recall/F1, where $TP/FP/FN$ are defined after one-to-one matching (at a chosen IoU threshold). This style of evaluation is used in biomedical instance challenges such as GlaS[204]:

$$\mathrm{Precision} = \frac{TP}{TP+FP}, \qquad \mathrm{Recall} = \frac{TP}{TP+FN}, \qquad \mathrm{F1} = \frac{2TP}{2TP+FP+FN}.$$

When methods output calibrated confidence scores, average precision (AP) is also informative. If $\mathrm{P}(r)$ denotes the precision as a function of recall $r \in [0,1]$, then

$$\mathrm{AP} = \int_0^1 \mathrm{P}(r)\, dr,$$

and the mean AP (mAP) averages AP across classes (and optionally across IoU thresholds).

**Object delineation quality (how good are the matched masks?).** Biomedical instance benchmarks frequently report object-level shape scores, especially when systems produce deterministic masks without calibrated confidence scores. Let $G_i$ be the $i$-th ground-truth instance and $S_j$ be the $j$-th predicted instance. Let $S_*(G_i)$ be the predicted instance with the largest overlap with $G_i$ (or an empty mask if none overlaps), and $G_*(S_j)$ the analogous ground-truth partner of $S_j$.

$$\mathrm{Dice}(A,B) = \frac{2|A \cap B|}{|A| + |B|}.$$

Then the symmetric object-level Dice is

$$D_{\mathrm{obj}}(G,S) = \frac{1}{2}\left(\frac{1}{n}\sum_{i=1}^{n} \mathrm{Dice}\big(G_i, S_*(G_i)\big) + \frac{1}{m}\sum_{j=1}^{m} \mathrm{Dice}\big(G_*(S_j), S_j\big)\right),$$

and an analogous symmetric object-level Hausdorff distance is

$$H_{\mathrm{obj}}(G,S) = \frac{1}{2}\left(\frac{1}{n}\sum_{i=1}^{n} \mathrm{HD}\big(G_i, S_*(G_i)\big) + \frac{1}{m}\sum_{j=1}^{m} \mathrm{HD}\big(G_*(S_j), S_j\big)\right).$$

(If unmatched instances are paired to an empty mask, Dice contributes 0; for distance metrics, a standard practice is to treat unmatched objects as failures and report them separately via $FP/FN$ and $F_1$.)

**Aggregated overlap metrics tailored to nuclei and small-object regimes.** For dense nuclei segmentation, many works and challenges use metrics designed to penalize splits/merges more directly than pixel IoU. A widely adopted example is the Aggregated Jaccard Index (AJI)[205] used in MoNuSeg evaluation[206]. Conceptually, AJI aggregates intersections and unions over best-overlap matches for each ground-truth instance and adds unmatched predictions as an explicit penalty term. Let $m(i)$ be the index of the predicted instance best matched to $G_i$ (or undefined if none matches), and let $U$ be the set of unmatched predicted instances. Then

$$\mathrm{AJI} = \frac{\sum_{i=1}^{n} |G_i \cap \hat{S}_{m(i)}|}{\sum_{i=1}^{n} |G_i \cup \hat{S}_{m(i)}| \;+\; \sum_{k \in U} |\hat{S}_k|}.$$

**Task-specific bundles for instance segmentation.** Because downstream use varies, we recommend choosing primary endpoints according to the dominant error mode:

**Sparse lesion detection** (for example liver or lung lesions): treat the task as detection dominated. Report lesion-wise sensitivity and false-positive rate per volume at clinically relevant size thresholds, and use AP or F1 at fixed IoU as secondary summaries.

**Dense nuclei or cell segmentation**: prioritize metrics that penalize splits and merges, such as object-level F1 at fixed IoU and AJI. Where possible, provide an error breakdown into missed, split, and merged instances.

**Count-critical applications** (for example glomeruli counts, micro-metastases): complement segmentation metrics with the absolute or relative error in per-case counts, because this is often the quantity used for clinical decision making.

**Recommended protocol for instance-level biomedical segmentation.** Across these settings, a minimal protocol is:

- Specify and fix the matching rule (IoU threshold, assignment algorithm) and report it alongside results.
- Report both detection quality (object-level precision, recall, F1 or AP) and delineation quality (object-level Dice or HD) rather than only pixel-wise overlap.
- For multi-class instances, provide class-wise scores and macro/micro averages, and avoid relying solely on a single global mean.
- When data are highly crowded or labels are noisy, include qualitative error analysis and per-case distributions to reveal systematic under- or over-segmentation.

##### 5.1.3 Panoptic Segmentation

In biomedical imaging, panoptic formulations are often motivated less by "scene parsing" in the computer-vision sense and more by the need to connect compartment-level morphology to cell-level composition. Digital pathology is a canonical example: clinically meaningful readouts such as tumor-immune interactions, stromal inflammation, and tumor-infiltrating lymphocyte (TIL) burden depend on simultaneously delineating tissue regions and identifying cell instances within those regions. The PUMA melanoma benchmark[207] explicitly operationalizes this two-level requirement by pairing tissue semantic segmentation with nuclei instance segmentation and reporting Dice for tissue regions and class-wise (micro and average) F1 for nuclei. Likewise, PanopTILs[208] is introduced specifically to enable joint region and cell modeling for explainable TIL scoring, and reports performance both at the segmentation level and at clinically aligned endpoints.

**Generic panoptic metrics.** When a task genuinely requires a single unified score across "stuff" (regions) and "things" (instances), the dominant metric is Panoptic Quality (PQ)[209], computed per class and then averaged. PQ matches predicted and ground-truth segments (commonly using an IoU threshold such as 0.5) and combines recognition and mask overlap into one quantity. For a given class $c$, after matching predicted segments $p$ with ground-truth segments $g$, define: $TP_c$: matched pairs $(p, g)$; $FP_c$: unmatched predictions; $FN_c$: unmatched ground truths.

$$\mathrm{PQ}_c = \frac{\sum_{(p,g)\in \mathrm{TP}_c} \mathrm{IoU}(p,g)}{|\mathrm{TP}_c| + \frac{1}{2}|\mathrm{FP}_c| + \frac{1}{2}|\mathrm{FN}_c|}.$$

Also:

$$\mathrm{SQ}_c = \frac{1}{|\mathrm{TP}_c|}\sum_{(p,g)\in \mathrm{TP}_c} \mathrm{IoU}(p,g), \quad \mathrm{RQ}_c = \frac{|\mathrm{TP}_c|}{|\mathrm{TP}_c| + \frac{1}{2}|\mathrm{FP}_c| + \frac{1}{2}|\mathrm{FN}_c|}, \quad \mathrm{PQ}_c = \mathrm{SQ}_c \cdot \mathrm{RQ}_c.$$

In biomedical reports, it is often informative to stratify PQ into $PQ_{\text{things}}$ (cells/lesions) and $PQ_{\text{stuff}}$ (anatomy/tissue compartments), because dominant failure modes differ (crowding and touching objects versus ambiguous region boundaries and stain variability).

**Biomedical "panoptic by construction" (region Dice + nuclei instance scores).** Many pathology benchmarks that call themselves "panoptic" in practice evaluate the two components with domain-preferred metrics: Dice for region masks and $F_1$-style criteria for cell instances. PUMA[207], for example, reports average and micro-average Dice over tissue classes and class-wise $F_1$, macro-$F_1$, and micro-$F_1$ for nuclei. This decomposition is clinically sensible because downstream readouts are typically computed by intersecting the two outputs; reporting each component separately makes error attribution and method comparison far clearer than a single scalar.

For a tissue class $k$ with predicted pixel set $P_k$ and ground-truth pixel set $G_k$:

$$\mathrm{Dice}_k = \frac{2|P_k \cap G_k|}{|P_k| + |G_k|} = \frac{2\mathrm{TP}_k}{2\mathrm{TP}_k + \mathrm{FP}_k + \mathrm{FN}_k}.$$

Micro-averaged Dice aggregates counts over classes (and optionally over images):

$$\mathrm{Dice}_{\text{micro}} = \frac{2\sum_k \mathrm{TP}_k}{2\sum_k \mathrm{TP}_k + \sum_k \mathrm{FP}_k + \sum_k \mathrm{FN}_k}.$$

For an instance class $c$ (e.g., nuclei type), after a chosen matching rule:

$$\mathrm{Precision}_c = \frac{\mathrm{TP}_c}{\mathrm{TP}_c + \mathrm{FP}_c} \qquad \mathrm{Recall}_c = \frac{\mathrm{TP}_c}{\mathrm{TP}_c + \mathrm{FN}_c} \qquad F_{1,c} = \frac{2\,\mathrm{Precision}_c\,\mathrm{Recall}_c}{\mathrm{Precision}_c + \mathrm{Recall}_c}.$$

Macro and micro averaging across $C$ classes:

$$F_{1,\text{macro}} = \frac{1}{C}\sum_{c=1}^{C} F_{1,c},$$

$$\mathrm{Precision}_{\text{micro}} = \frac{\sum_c \mathrm{TP}_c}{\sum_c(\mathrm{TP}_c + \mathrm{FP}_c)} \qquad \mathrm{Recall}_{\text{micro}} = \frac{\sum_c \mathrm{TP}_c}{\sum_c(\mathrm{TP}_c + \mathrm{FN}_c)} \qquad F_{1,\text{micro}} = \frac{2\,\mathrm{Precision}_{\text{micro}}\,\mathrm{Recall}_{\text{micro}}}{\mathrm{Precision}_{\text{micro}} + \mathrm{Recall}_{\text{micro}}}.$$

**Recommended protocol for panoptic biomedical segmentation.** When defining a panoptic benchmark or reporting results, we recommend:

- Decide whether the primary goal is a unified scene score or a decomposition into interpretable components. For the former, report per-class PQ, $PQ_{\text{things}}$, and $PQ_{\text{stuff}}$, together with SQ and RQ to separate recognition and mask quality.
- For region-plus-cell settings such as digital pathology, treat tissue regions and cell instances as distinct outputs. Report Dice (and HD95 where relevant) for tissue regions, and class-wise F1 or AP for cells, along with macro and micro averages.
- Where downstream endpoints (for example TIL burden, immune-hot/cold stratification) are available, include at least one analysis that relates segmentation performance to error in these endpoints, clarifying how much segmentation improvement translates into clinically meaningful gains.

#### 5.2 Federated learning (FL) and multi-site evaluation metrics

In FL, evaluation must quantify not only mean performance but also site-wise variability, because biomedical domain shift across institutions is often the main driver of real-world failure. A minimal reporting standard is: (i) per-site Dice/IoU (and distance metrics when relevant), plus (ii) dispersion summaries (standard deviation, worst-site performance, and percentiles).

Let there be $S$ sites. Let $M_s$ denote a metric (e.g., Dice) computed on site $s$. Then:

$$\overline{M} = \frac{1}{S}\sum_{s=1}^{S} M_s, \qquad \text{Std}(M) = \sqrt{\frac{1}{S-1}\sum_{s=1}^{S}(M_s - \overline{M})^2}, \qquad M_{\min} = \min_{1\le s\le S} M_s.$$

A simple heterogeneity gap is the worst-to-mean drop:

$$\Delta_{\text{worst}} = \overline{M} - M_{\min}.$$

Beyond segmentation accuracy, FL papers frequently use vector similarity metrics to quantify client contribution, update consistency, or agreement between client gradients/representations[210]. For example, client contribution estimation has used Pearson correlation, cosine similarity, and Euclidean distance between client update vectors. Let $u, v \in \mathbb{R}^d$ be two client update vectors (e.g., flattened gradients or parameter deltas).

Pearson correlation:

$$\rho(u,v) = \frac{\sum_{i=1}^{d}(u_i - \bar{u})(v_i - \bar{v})}{\sqrt{\sum_{i=1}^{d}(u_i - \bar{u})^2}\sqrt{\sum_{i=1}^{d}(v_i - \bar{v})^2}}.$$

Cosine similarity:

$$\text{CosSim}(u,v) = \frac{u^\top v}{\|u\|_2\,\|v\|_2}.$$

Euclidean distance:

$$\text{Dist}_2(u,v) = \|u - v\|_2 = \sqrt{\sum_{i=1}^{d}(u_i - v_i)^2}.$$

These metrics are not substitutes for downstream segmentation performance; rather, they are diagnostic signals for heterogeneity and optimization dynamics, and should be reported alongside (not instead of) site-wise segmentation outcomes[210].

**Recommended protocol for multi-site and FL evaluation.** To make robustness to domain shift explicit, we suggest the following minimum protocol:

- Always report per-site segmentation metrics (for example Dice and HD95 for key structures) rather than only a pooled mean over all sites.
- Provide dispersion summaries across sites, including standard deviation, interquartile range, and worst-site performance; include the worst-to-mean gap $\Delta_{\text{worst}}$ as a simple heterogeneity indicator.
- When sites differ systematically (for example scanner vendor, acquisition protocol, population), stratify performance by these factors and highlight any systematic biases.
- If the model is intended for deployment across sites, define task-specific safety thresholds (for example per-site Dice $> 0.8$ for critical structures) and report how many sites and which ones fall below these thresholds.
- Use gradient or update similarity metrics only as auxiliary diagnostics; they should complement, not replace, site-wise segmentation outcomes.

#### 5.3 Prompt-based and Interactive Segmentation Metrics

Promptable foundation models change what "good" means: in many biomedical workflows, the scarce resource is not FLOPs but expert time. Evaluation should therefore quantify quality as a function of interaction and explicitly report the cost needed to reach acceptable contours.

**Quality-vs-interaction curves and area-under-curve summaries.** Let $Q_i(k)$ be a quality metric for case $i$ after $k$ interactions (clicks, scribbles, corrective prompts). A standard summary is the mean interaction curve $\overline{Q}_k$ and its area under curve up to a maximum budget $K$. This is used directly in interactive medical benchmarks, including DSC-AUC and NSD-AUC reporting[211].

$$\bar{Q}(k) = \frac{1}{N}\sum_{i=1}^{N} Q_i(k), \qquad \mathrm{AUC}(Q) = \frac{1}{K+1}\sum_{k=0}^{K} \bar{Q}(k).$$

This formulation makes two biomedical-relevant comparisons explicit: (i) which method gives higher quality under small interaction budgets (typical clinical time constraints), and (ii) which method saturates fastest.

**Time-aware summaries (quality as a function of annotation time).** If $\Delta t_i(\ell)$ is the time spent on the $\ell$-th interaction for case $i$, define cumulative time $T_i(k) = \sum_{\ell=1}^{k} \Delta t_i(\ell)$ and a time-parameterized curve $Q_i(T)$. A practical summary is the quality achieved under a fixed time budget $T_{\max}$:

$$Q_i(T_{\max}) = \max\{Q_i(k) : T_i(k) \le T_{\max}\}, \qquad \overline{Q}(T_{\max}) = \frac{1}{N}\sum_{i=1}^{N} Q_i(T_{\max}).$$

(Reporting both click budgets and time budgets is recommended when interaction primitives differ across methods.)

**Number of clicks to reach a target threshold (NoC) and failure rate.** A clinically interpretable metric is the Number of Clicks (NoC) required to achieve a predefined threshold, for example DSC $\ge 0.85$ or HD95 $\le 5$ mm. Recent interactive tumor-contouring studies report NoC for multiple DSC/HD95 thresholds, together with a capped interaction budget and an explicit failure rate[212].

$$\mathrm{NoC}_i(\tau) = \min\{k \in \{0, \dots, K_{\max}\} : \; Q_i(k) \ge \tau\}, \qquad \mathrm{NoC}(\tau) = \frac{1}{N}\sum_{i=1}^{N} \mathrm{NoC}_i(\tau).$$

If a case fails to reach the target within $K_{\max}$, it is counted as a failure, and the Percent of Failures (PoF) is reported:

$$\mathrm{PoF} = \frac{\#\{i : \; \mathrm{NoC}_i(\tau) \text{ undefined within } K_{\max}\}}{N} \times 100\%.$$

This pair (NoC, PoF) captures "how often will the expert be forced to give up or switch tools," which is often more operationally meaningful than marginal Dice gains.

**Practical protocol for interactive and prompt-based evaluation.** To make interaction efficiency and robustness comparable across methods, we recommend:

- Fix a maximum interaction budget $K_{\max}$ that approximates realistic clinical constraints (for example a small number of clicks per case), and use the same budget for all methods.
- Standardize interaction primitives and policies: define how positive and negative clicks are placed (for example always at the largest remaining error region) and whether they are provided by humans or simulated annotators.
- Report both DSC-AUC or NSD-AUC over $k = 0, \dots, K_{\max}$ and NoC/PoF at clinically meaningful quality thresholds (for example DSC $\ge 0.85$ and NSD $\ge 0.9$), making clear how often the model fails to reach the target.
- When time measurements are available, complement click-based summaries with $Q(T_{\max})$ under realistic time budgets (for example 30–60 seconds per case), since different interaction modalities can have different time costs.
- For prompt-only, non-interactive uses of foundation models, explicitly state the prompt design, any test-time prompt optimization, and whether reported metrics correspond to a single prompt, per-case tuned prompts, or an oracle setting.

Taken together, these elements align evaluation with the practical question faced by clinicians: how much expert effort is required to obtain contours that are good enough for the downstream task.
